# Division-Specific Organization of a Shared Functional Scaffold in the Early-Life Human Brain

**DOI:** 10.64898/2026.09.14.751555

**Authors:** Dan Hu, Jiale Cheng, Kangfu Han, Zhengwang Wu, Weiyan Yin, Yue Sun, Jiameng Liu, Sheng-Che Hung, Li Wang, Jessica R. Cohen, Weili Lin, Gang Li, UNC/UMN Baby Connectome Project Consortium

## Abstract

Early cognition and behavior emerge from coordinated maturation of functional systems spanning the brain, making cross-division integration essential for defining whole-brain architecture and understanding its role in neurodevelopment. Yet early human functional development is still largely investigated through cortical networks or through isolated, coarsely resolved noncortical structures, leaving unclear how this integrated architecture is organized across the cerebral cortex, subcortex and cerebellum. Moving beyond structure-isolated and adult-derived maps, we create a reproducible, fine-grained Early-Life Whole-Brain Functional Parcellation (UNC-ELF) spanning birth to six years. Using this framework, we identify a shared functional scaffold expressed through distinct division-specific modes: differentiated and dual-polarity cortical organization, spatially graded cerebellar organization with blended functional transition zones, and nucleus-constrained subcortical mosaics of multi-system affiliation. Across early childhood, the scaffold is selectively refined through heterogeneous, nonlinear connectivity trajectories, with major inflections concentrated in infancy. Age-, sex- and prospective cognition-related variation is differentially encoded across distinct connectomic components. Together, these findings reframe early functional brain development as coordinated refinement of a shared but nonuniform whole-brain scaffold and establish UNC-ELF as a unified reference for resolving typical and atypical functional organization across the developing brain.

## INTRODUCTION

Early childhood is a period of rapid functional reorganization during which large-scale brain systems emerge, differentiate and establish the foundations for later cognition and behavior^1–3^. This development does not occur within the cerebral cortex alone, but depends on coordinated maturation across the cerebral cortex, subcortex, and cerebellum. Functional organization is already detectable across these divisions near birth: thalamocortical connectivity aligns with emerging cortical systems, while the cerebellum exhibits organized topography and increasingly differentiated cortical interactions across early childhood^4,5^. Yet it remains unclear whether these observations reflect largely independent developmental processes or the division-specific expression of a shared whole-brain functional architecture. Resolving this question requires a common developmental reference capable of characterizing how functional systems are organized and coordinated across the major brain divisions.

Existing functional parcellations have provided essential reference systems for studying large-scale brain organization, but their suitability for resolving coordinated early development remains limited. Most widely used parcellations were derived from mature brains, often under different anatomical assumptions and at distinct developmental stages^6–9^. This age mismatch is consequential, as mature brains exhibit relatively stabilized and segregated functional topography, whereas early development is characterized by more dynamic and transitional architecture. Neurodevelopment-specific parcellations have begun to address this gap, including fine-grained maps of the infant cerebral cortex^10,11^ and early functional mapping of the cerebellum^5^. Yet these advances remain largely structure-specific. Major brain divisions are typically parcellated separately, often from different cohorts and with distinct preprocessing and analytical strategies. Consequently, parcel boundaries, network labels, and organizational scales are not directly aligned across structures, and combining these independent maps can introduce methodological confounds. This challenge is particularly relevant for the subcortex and cerebellum. Many noncortical mapping approaches assign territories to predefined cortical networks according to their strongest functional connectivity^8,12–14^, thereby imposing a cortical reference that may not fully capture developmentally evolving cross-structure relationships. Conversely, data-driven developmental efforts often define noncortical parcels using only within-structure connectivity^5,10,11^, potentially overlooking cerebello-cortical and subcortico-cortical interactions that contribute to functional boundary organization. These limitations make it difficult to determine whether apparent cross-structure differences reflect genuine division-specific neurobiology or differences introduced by parcellation design, leaving early whole-brain functional architecture without a unified developmental reference.

Here, we address this gap by constructing an Early-Life Whole-Brain Functional Parcellation (UNC-ELF) spanning birth to six years within a common developmental framework. Parcellations of the cerebral cortex, cerebellum and subcortex were derived from the same early-life cohorts using a shared analytical strategy with structure-adapted connectivity features. Cortical parcels were defined from intrinsic cerebral connectivity, whereas cerebellar and subcortical parcels incorporated both within-division and cortex-centered connectivity, allowing local organization and broader cross-structure relationships to be represented within the same reference frame. Rather than combining independently derived maps, this framework enables direct comparison of functional correspondence across the major brain divisions.

Using this framework, we show that early-life functional organization is structured across the whole brain but is not expressed as a uniform replication of cortical networks. Instead, a shared cortex-referenced scaffold is manifested through distinct division-specific modes: differentiated organization in cortex, spatially graded cerebellar organization and locally heterogeneous, nucleus-constrained mosaics in subcortex. We further establish the reproducibility, developmental appropriateness and external generalizability of the parcellation and use it to characterize how whole-brain connectivity is selectively refined through division- and connection-specific developmental trajectories. Predictive analyses further indicate that developmental and cognitive information is distributed across distinct components of the scaffold, with cortical, subcortical and cerebellar connectivity contributing differentially across outcomes. Together, these findings establish a unified developmental reference for resolving how a shared whole-brain functional scaffold is differentially organized, refined and functionally expressed across the major brain divisions.

## RESULTS

### A unified framework identifies a shared early-life whole-brain functional scaffold

To determine whether early functional brain organization is coordinated across the cerebral cortex, subcortex and cerebellum, we constructed the Early-Life Whole-Brain Functional Parcellation (UNC-ELF), spanning birth to six years. UNC-ELF resolved the three major brain divisions within a unified developmental framework while preserving structure-adapted connectivity features. Cortical parcels were defined from intrinsic cerebral connectivity, whereas cerebellar and subcortical parcels were informed by both within-structure and cortex-centered connectivity. The principal solution comprised 22 cerebral, 29 cerebellar and 31 subcortical networks across six major bilateral nuclei (Fig. 1a). Higher-resolution 36-network cerebral and 54-network cerebellar solutions are provided in Extended Data Fig. 2. Across divisions, UNC-ELF exhibited pronounced bilateral symmetry and anatomically coherent spatial organization, providing a structured reference for identifying shared and division-specific features of early whole-brain functional architecture.

**Figure 1.**
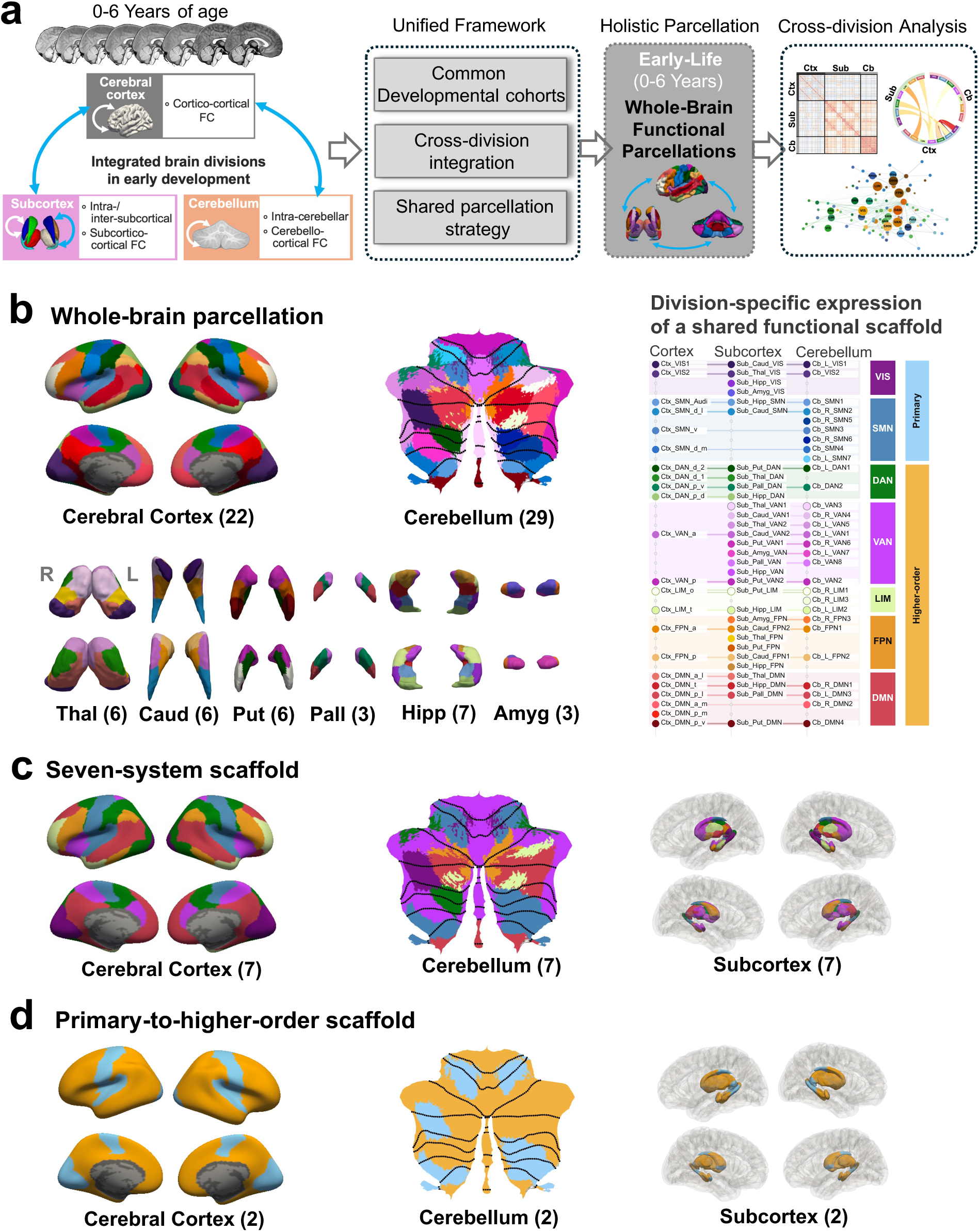
Unified early-life whole-brain functional parcellation reveals a shared cortex-referenced scaffold with division-specific expression. **a,** Schematic of the unified whole-brain parcellation framework. Resting-state fMRI data acquired from birth to 6 years were analyzed within a common developmental framework to define functional territories across the cerebral cortex, subcortex and cerebellum. A shared parcellation strategy combined division-specific intrinsic connectivity with cross-division connectivity features to generate the Early-Life Whole-Brain Functional Parcellation (UNC-ELF) and establish a cortex-referenced framework for cross-structure correspondence. **b,** Principal parcellation and fine-scale cross-division correspondence. The solution comprises 22 cortical networks, 29 cerebellar networks and 31 subcortical parcels distributed across the thalamus (Thal, 6), caudate (Caud, 6), putamen (Put, 6), pallidum (Pall, 3), hippocampus (Hipp, 7) and amygdala (Amyg, 3). Cortical networks are displayed on the cerebral surface, cerebellar networks on the SUIT cerebellar flatmap and subcortical parcels as nucleus-specific surface renderings. Shared hues indicate correspondence to the same cortex-referenced functional family across divisions. The division-specific scaffold map aligns corresponding cortical, subcortical and cerebellar parcels within the seven large-scale systems and further distinguishes their expression within primary and higher-order functional classes, revealing both shared functional identity and division-specific organization. **c,** Collapse of the fine-grained parcellation into seven large-scale functional systems, revealing a shared system-level scaffold across the cerebral cortex, subcortex and cerebellum. **d,** Collapse into primary and higher-order classes, showing the broad functional hierarchy represented across all three divisions. Network abbreviations: VIS, visual; SMN, somatomotor; DAN, dorsal attention; VAN, ventral attention; LIM, limbic; FPN, frontoparietal; DMN, default mode. Brain division abbreviations: Ctx, cerebral cortex; Sub, subcortex; Cb, Cerebellum.

The cerebral cortex was chosen as the common reference backbone because its highly differentiated large-scale system architecture was evident near birth and remained broadly conserved across age-specific solutions, providing a stable developmental coordinate system for relating the cerebellum and subcortex (Extended Data Fig. 5). We then grouped the data-driven 22-network cortical solution into seven large-scale systems corresponding to canonical functional topology and assigned each cerebellar and subcortical parcel according to its cortex-centered connectivity profile (Fig. 1c). This strategy linked locally derived cerebellar and subcortical territories to a common cortical reference while allowing each division to retain its characteristic spatial organization. All seven systems were represented in both noncortical divisions, revealing a shared system-level architecture extending beyond the cortex. In the cerebellum, somatomotor territories were concentrated in the anterior lobe and lobule VIII, whereas default-mode and frontoparietal territories extended across the posterior lobe, particularly Crus I and Crus II. In the subcortex, the same systems were expressed as heterogeneous, nucleus-constrained mosaics: higher-order correspondences were broadly represented in the thalamus, caudate and hippocampus, whereas sensorimotor correspondence was most prominent in the putamen and selected thalamic territories. Collapsing these assignments into primary and higher-order classes (Fig. 1d) further showed that the cortical functional hierarchy was broadly represented beyond the cortex, establishing a shared whole-brain scaffold expressed through division-specific topographies.

We next asked whether the shared seven-system scaffold could be resolved into finer cross-division functional families. Within each large-scale system, cerebellar and subcortical parcels were matched to specific fine-scale networks of the 22-network cortical solution according to their reproducible cortex-centered connectivity profiles. The resulting division-specific scaffold map aligns corresponding cortical, subcortical and cerebellar parcels within each of the seven systems, while also distinguishing primary from higher-order functional classes (Fig. 1b). Under a conservative matching criterion permitting at most one cerebellar and one subcortical parcel per cortical network, 15 of the 22 cortical networks showed three-way correspondence across all divisions. These included every visual, ventral-attention, limbic and frontoparietal family, together with selected somatomotor, dorsal-attention and default-mode families. Thus, the broad seven-system scaffold was further resolved into a nested set of cross-division functional correspondences, revealing a whole-brain architecture in which shared functional identities are preserved across structures but expressed through division-specific spatial organization.

### The shared scaffold is expressed through distinct division-specific organizational modes

The shared scaffold, defined across seven large-scale systems and finer cortex-anchored functional correspondences, captured the dominant affiliations of parcels across brain divisions but concealed marked variation in the specificity of those affiliations. Cortical networks formed comparatively differentiated territories, whereas cerebellar and subcortical parcels ranged from highly specific cortical-system counterparts to reproducible multi-system affiliations with region-specific stability and separability. To resolve this variation, we jointly examined assignment stability, separability between the two strongest cortical correspondences and secondary system admixture (Figs. 2 and 3) across bootstrap resamples. High stability together with high separability identified well-defined cortical correspondence, whereas high stability with low separability indicated reproducible convergence of multiple system affiliations rather than unstable assignment.

**Figure 2.**
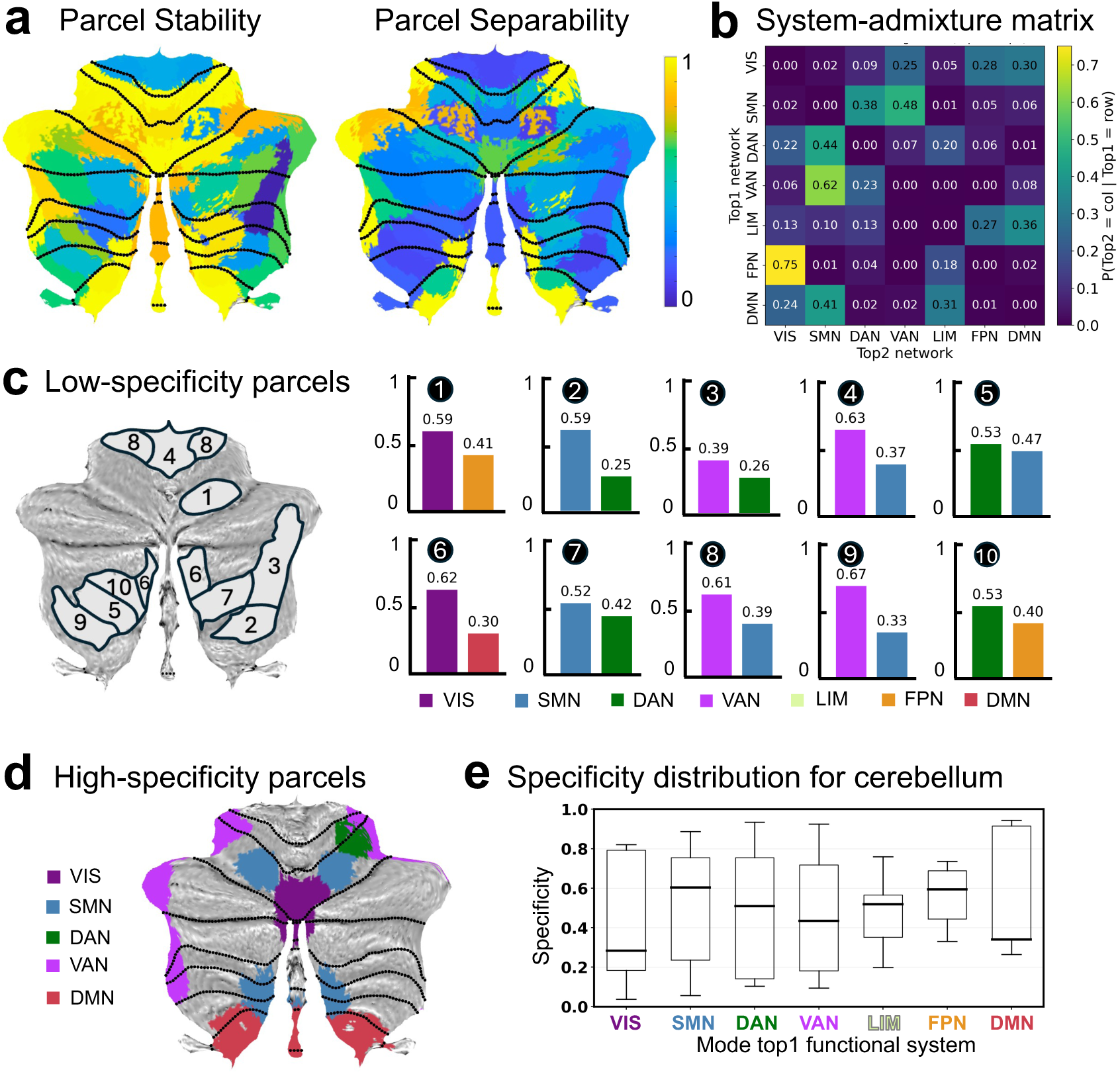
The cerebellum exhibits spatially graded correspondence to cortical functional systems. Analyses were performed using the 54-network cerebellar solution, providing a finer spatial scale for resolving cerebello-cortical correspondence and local functional admixture. **a**, Cerebellar flatmaps showing parcel stability and separability across bootstrap-based assignments to cortical systems. Stability denotes the probability of the dominant cortical-system assignment across bootstrap iterations. Separability denotes the median margin between the strongest and second-strongest correspondence scores across bootstraps, with higher values indicating more distinct cortical-system correspondence. Composite specificity was defined from the joint consideration of stability and separability. **b**, System-admixture matrix summarizing secondary cortical-system affiliations. Entries indicate the probability that a secondary system assignment (Top2; columns) occurs given a dominant system assignment (Top1; rows). Off-diagonal values indicate structured mixture between cortical systems rather than diffuse assignment ambiguity. **c**, Representative low-specificity cerebellar parcels and their two leading cortical-system assignment probabilities. Black circled numbers indicate rank order among the displayed low-specificity parcels, with 1 denoting the lowest specificity score. Bars show the two systems most frequently assigned across bootstrap iterations, with values indicating their assignment probabilities. Low-specificity parcels were concentrated near interfaces between functional territories and exhibited reproducible blended multi-system affiliations. **d**, Spatial distribution of high-specificity cerebellar parcels, defined by jointly high stability and separability. These parcels formed spatially coherent functional cores with well-differentiated cortical-system correspondence. **e**, Distribution of composite specificity across cerebellar parcels grouped by their modal cortical-system assignment. Broad within-system variation reflects a continuum from high-specificity cores to lower-specificity transition zones. Of note, because SUIT flatmaps project three-dimensional cerebellar labels onto a two-dimensional surface using mode-based assignment, visually adjacent regions in **c** and **d** may correspond to distinct volumetric parcels. For box plots: center line, median; box limits, upper and lower quartiles; whiskers, 1.5× interquartile range.

**Figure 3.**
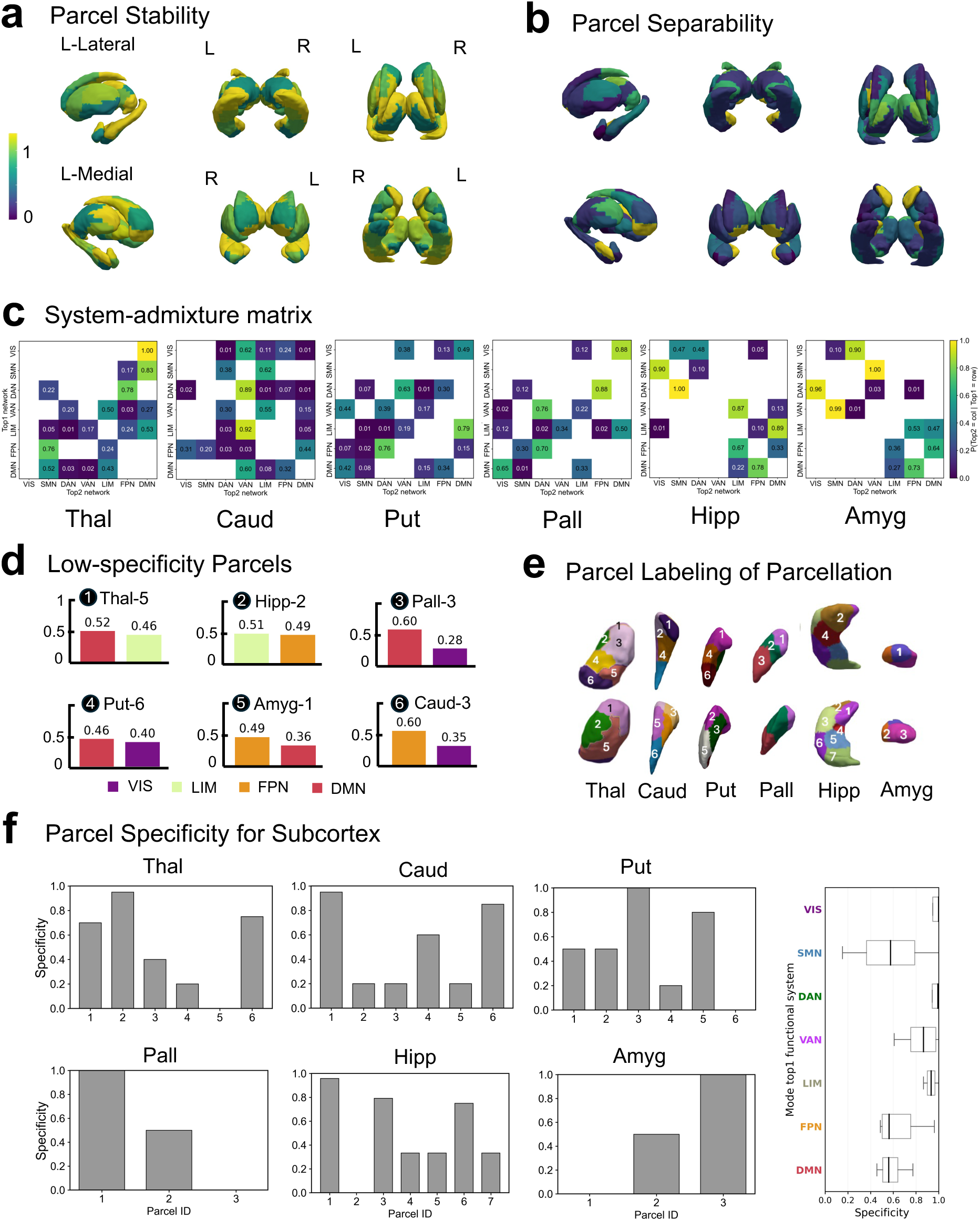
The subcortex forms nucleus-constrained functional mosaics with heterogeneous multi-system affiliations. **a**, Parcel stability mapped onto subcortical structures. Stability denotes the bootstrap probability of the modal cortical-system assignment, with higher values indicating more reproducible cortex-referenced correspondence. **b**, Spatial distribution of parcel separability, defined as the median margin between the strongest and second-strongest cortical-system correspondence scores across bootstrap iterations. Higher values indicate more distinct system affiliation. **c**, Nucleus-specific system-admixture matrices. Entries show the probability of a secondary system correspondence (Top2; columns) conditional on the dominant correspondence (Top1; rows). Off-diagonal values reveal structured mixture among cortical-system affiliations within each nucleus; values below 0.005 were masked for visualization. **d**, Representative low-specificity subcortical parcels. Specificity summarizes both stability and separability; lower values indicate less distinct or more admixed cortical-system correspondence. Black circled numbers indicate rank order among the displayed low-specificity parcels after ranking them together across subcortical nuclei, with 1 denoting the lowest specificity score. Parcel labels indicate the host nucleus and parcel ID shown in e. Bars show the two systems most frequently assigned across bootstrap iterations, with values indicating their assignment probabilities. These parcels show mixed multi-system affiliations, predominantly exhibiting secondary affiliations with higher-order default-mode and frontoparietal networks. **e**, Anatomical rendering of the subcortical parcellation with parcel IDs corresponding to the labels used in d and f. **f**, Parcel- and system-level specificity across subcortical nuclei. Bar plots show the composite specificity score for each parcel within the thalamus, caudate, putamen, pallidum, hippocampus and amygdala, computed as the average of percentile-ranked stability and percentile-ranked separability. The coexistence of high- and low-specificity parcels within individual nuclei highlights the internally heterogeneous, mosaic-like organization of the subcortex. The boxplot summarizes dominant-system assignment specificity across all subcortical parcels, grouped by their assigned cortical functional system. Assignment specificity is defined as the bootstrap probability of the assigned Top1 system. Visual, dorsal-attention, ventral-attention and limbic parcels generally show strongly dominant assignments, whereas somatomotor, frontoparietal and default-mode parcels exhibit broader distributions and greater parcel-level heterogeneity. Boxes indicate the interquartile range, center lines indicate medians and whiskers extend to 1.5 times the interquartile range.

In the cerebellum, assignment stability was broadly high, while cortical correspondence followed a spatially graded core–transition organization (Fig. 2a, e). High-specificity parcels, defined by jointly high stability and high separability, formed segregated functional cores across both primary and association zones (Fig. 2d), whereas low-specificity parcels clustered near interfaces between these cores (for example, near the lobule VI-Crus I interface; Fig. 2c). Their secondary affiliations were structured rather than diffuse: somatomotor-aligned territories most often showed additional correspondence with ventral- and dorsal-attention systems, whereas visual territories showed secondary correspondence with default-mode, frontoparietal and ventral-attention systems (Fig. 2b). More broadly, cerebellar admixture was dominated by primary-to-higher-order overlap, particularly along somatomotor-attention and visual-association axes. These patterns indicate a graded cerebellar organization in which stable functional cores are bordered by blended transition zones with reproducible multi-system affiliation.

The subcortex exhibited a distinct organizational mode in which cortical-system correspondence was locally heterogeneous but constrained by nuclear anatomy (Fig. 3). When system correspondence was evaluated separately within each subcortical structure, canonical cortical systems were not cleanly segregated by nuclear boundaries. Instead, each nucleus contained multiple cortical-system correspondences arranged as an internal mosaic (Fig. 3e). High-specificity parcels, particularly those aligned with visual, dorsal-attention, ventral-attention and limbic systems, were spatially concentrated within individual nuclei, including selected hippocampal and striatal subdomains, and showed relatively well-defined cortico-subcortical correspondence (Fig. 3e, f). By contrast, low-specificity parcels were more often affiliated with default-mode and frontoparietal systems, suggesting greater network admixture within higher-order cortico-subcortical correspondence (Fig. 3d). This admixture was strongly shaped by host nucleus (Fig. 3c). Thalamus and pallidum showed prominent primary-to-association mixing, including visual-to-default-mode correspondence in thalamus and dorsal-attention-to-frontoparietal correspondence in pallidum. Caudate and putamen were dominated by attention-, limbic- and sensorimotor-related overlap, including limbic-to-ventral-attention mixing in caudate and frontoparietal-to-dorsal-attention mixing in putamen. Hippocampus and amygdala showed a two-block admixture structure, separating visual-sensorimotor-attention correspondences from limbic-frontoparietal-default-mode correspondences, exemplified by dorsal-attention-to-somatomotor mixing in hippocampus and somatomotor-to-ventral-attention mixing in amygdala. Subcortical admixture was therefore not globally diffuse, but organized as nucleus-constrained mosaics of specific and mixed multi-system affiliation.

Together, these analyses identify three organizational modes within the shared early-life scaffold: differentiated network territories in cortex, spatially graded core-transition organization in cerebellum and locally heterogeneous, nucleus-constrained mosaics in subcortex. The scaffold is shared at the level of functional identity, but its expression is not uniformly categorical. Instead, cortical affiliations vary in specificity and converge reproducibly within individual noncortical territories.

### Cross-structure connectivity supports a reproducible and developmentally appropriate whole-brain parcellation

The division-specific correspondence patterns suggested that noncortical boundaries may be shaped not only by local connectivity but also by interactions with the cerebral cortex. We therefore tested whether incorporating cross-structure functional connectivity improved boundary definition relative to parcellations derived from within-structure connectivity alone. Boundary contrast, defined as the difference between within-parcel and cross-boundary similarity of cross-structure connectivity profiles, was consistently higher for the UNC-ELF parcellations across BCP, Calgary and Pixar datasets. Improvements were significant in the cerebellum for all three datasets (all FDR-corrected P ≤ 5.6 × 10^-4^) and in the subcortex for Calgary and Pixar (all FDR-corrected P ≤ 1.1 × 10^-6^), across both 22- and 36-network cortical targets. Thus, cross-structure connectivity contributed information beyond local connectivity alone, sharpening functionally meaningful cerebellar and subcortical boundaries across independent datasets.

We next evaluated internal reproducibility across the age-independent and age-dependent solutions with test-retest data partitions (Fig. 4). Age-dependent and age-independent solutions showed closely matched Dice and silhouette coefficients across cerebral cortex, cerebellum and subcortex, indicating that the recovered organization reflected a stable developmental scaffold rather than split-specific clustering. These analyses supported two robust resolutions in the cerebral cortex (k = 22 and 37) and cerebellum (k = 29 and 54), whereas optimal subcortical resolution varied by nucleus: six networks each for the thalamus, caudate and putamen; three networks each for the pallidum and amygdala; and seven networks for the hippocampus (Fig. 4a-c). Parcellations derived independently from test and retest datasets were highly concordant, with subcortical Dice coefficients approaching 1.0. Spatial renderings similarly showed close recovery of cortical and cerebellar territories across partitions (Fig. 4a, b) and of parcel boundaries within each subcortical nucleus (Extended Data Fig. 3).

**Figure 4.**
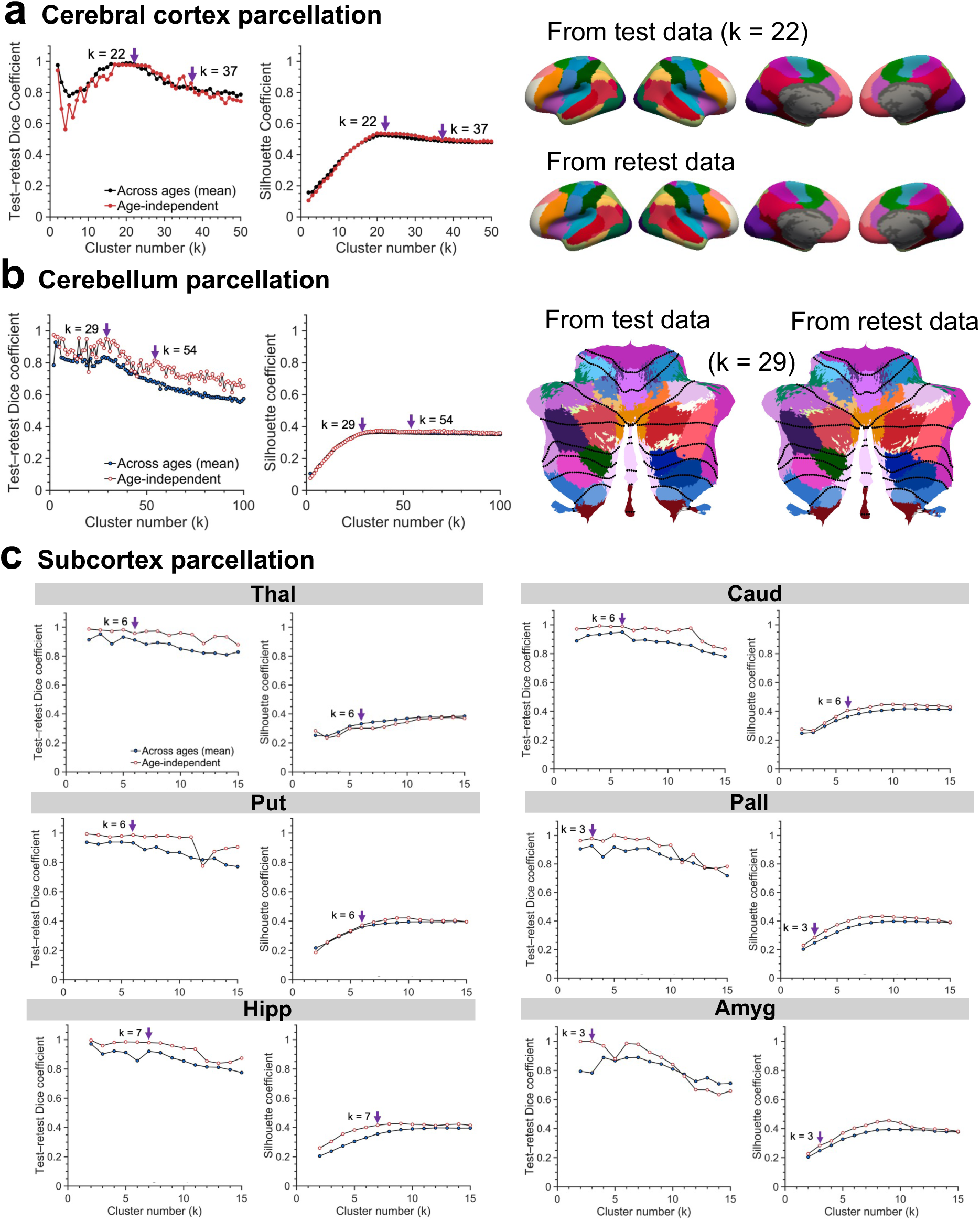
UNC-ELF is highly reproducible across data partitions and developmental groupings. **a**, Test-retest Dice and silhouette coefficients across candidate cluster numbers (k) for the cerebral cortex. Black curves summarize age-dependent solutions averaged across developmental age groups, and red curves show the age-independent solution. Arrows indicate reproducible candidate resolutions at k = 22 and k = 37. Surface renderings show the close spatial correspondence of the principal 22-network solution derived independently from test and retest data. **b**, Corresponding test-retest Dice and silhouette coefficients for the cerebellum. Reproducibility profiles identify reproducible resolutions at k = 29 and k = 54. Flatmaps show the spatial correspondence of the principal 29-network solution across test and retest data, confirming stable functional segregation. **c**, Nucleus-specific test-retest Dice and silhouette coefficients across candidate cluster numbers for the thalamus, caudate, putamen, pallidum, hippocampus and amygdala. Arrows indicate the selected resolutions: k = 6 for the thalamus, caudate and putamen; k = 3 for the pallidum and amygdala; and k = 7 for the hippocampus. Test–retest Dice coefficients were high at the selected resolutions, indicating reproducible recovery of subcortical parcel boundaries. Corresponding test and retest renderings are shown in Extended Data Fig. 3.

Age-resolved analyses further confirmed the high reproducibility and reflected a broadly preserved developmental scaffold rather than repeated reconfiguration across age groups. At the principal resolutions, the 22-network cerebral, 29-network cerebellar and nucleus-specific subcortical solutions retained their overall macroscale scale organization from birth through six years, with developmental variation expressed primarily as local boundary refinements (Extended Data Figs. 5 and 6). The 36-network cerebral solution, derived from the 37-network solution to preserve cross-hemispheric symmetry, and the 54-network cerebellar solution were similarly conserved across age groups (Supplementary Fig. 2), indicating that early development primarily refines an already recognizable large-scale scaffold rather than replacing it with successive age-specific architectures.

We then tested whether this unified framework better captured early-life functional organization than commonly used adult-derived parcellations (Fig. 5). Across all brain divisions, the UNC-ELF parcellations showed greater within-network functional homogeneity at comparable spatial scales. In the cerebral cortex, collapsing the 22-network solution to 17 and 7 systems yielded higher homogeneity than the corresponding Yeo-17 and Yeo-7 references^6^ (*t* = 142.05, *P* < 10^−300^, and *t* = 47.64, *P* = 1.64 × 10^−209^, respectively). Moreover, overlap between UNC-ELF and adult cortical maps declined along the functional hierarchy (*ρ* = −0.82, *P* = 0.034), with closer correspondence in sensory systems and greater divergence in higher-order association networks. Adult-derived topographies therefore showed reduced correspondence in systems undergoing greater developmental reorganization.

**Figure 5.**
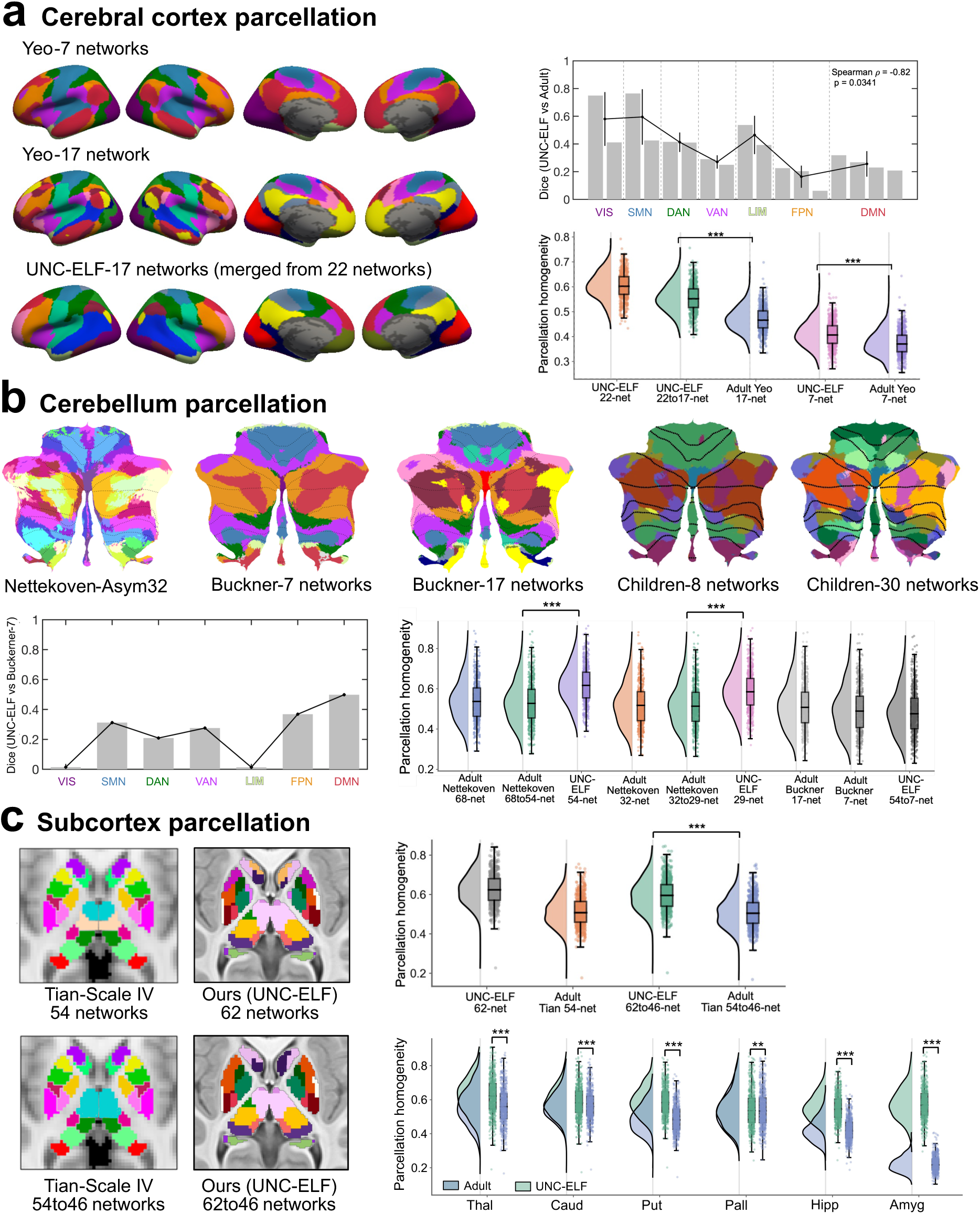
The UNC-ELF Parcellation better captures early developmental organization than adult-derived references. **a**, Comparison of the UNC-ELF cerebral cortical parcellation with adult-derived Yeo 17- and 7-network maps. Cortical surface renderings show the adult references and corresponding early-life representations obtained by merging the 22-network solution. Dice overlap between the UNC-ELF and adult 17-network maps decreased along the cortical functional hierarchy, with greatest divergence in higher-order frontoparietal (FPN) and default-mode (DMN) systems (Spearman *ρ* = −0.82, *p* = 0.034). Functional homogeneity was higher for the UNC-ELF cortical representations than for the adult references at both matched scales. **b**, Comparison of the UNC-ELF cerebellar solutions with adult-derived Nettekoven and Buckner parcellations. The 54- and 29-network UNC-ELF solutions showed higher functional homogeneity than resolution-matched Nettekoven maps. The seven-system UNC-ELF representation showed comparable homogeneity to the Buckner seven-network map while retaining cortex-referenced system correspondence. **c**, Comparison of the UNC-ELF subcortical parcellation with the adult-derived Tian Scale IV atlas after matching both schemes to a common 46-label resolution. Functional homogeneity was higher for the UNC-ELF parcellation across the whole subcortex and within each major nucleus. Violin plots show subject-level distributions; embedded boxes indicate the interquartile range, center lines the median and whiskers 1.5 times the interquartile range. Points denote individual participants. Statistical comparisons used one-sided paired *t*-tests. *** indicates *p* < 0.0001, ** indicates *p* < 0.01.

This developmental advantage extended beyond the cortex. Both the 54- and 29-network cerebellar solutions showed higher functional homogeneity than resolution-matched adult-derived comparators based on the Nettekoven parcellation^15^ (*t* = 82.70 and 70.55, respectively; both *P* < 10^−100^). The coarse seven-system cerebellar representation did not significantly exceed the Buckner seven-network solution^8^ in local homogeneity, consistent with its emphasis on cross-structure alignment rather than local compactness alone. In the subcortex, matching the UNC-ELF and the adult-derived Tian Scale IV parcellation^16^ to a shared 46-label resolution revealed markedly higher whole-subcortex homogeneity for the UNC-ELF solution (*t* = 237.30, *P* < 10^−300^), with the same advantage observed within each major nucleus. Qualitative comparison with a pediatric cerebellar parcellation^5^ showed broadly consistent anterior-posterior organization, with the present parcellation showing visually more compact and anatomically more regular parcels in lateral Crus I/II and posterior-inferior cerebellar regions (Fig. 5b).

The functional advantage of the UNC-ELF parcellation generalized to the independent Pixar^17^, Calgary^18^, and dHCP^19^, which span different developmental periods, acquisition protocols and behavioral states (Extended Data Fig. 4). Orthogonal validation using cortical morphology and tractography likewise showed greater within-parcel coherence than adult-derived reference schemes (Extended Data Fig. 8). Together, these findings show that cross-structure connectivity supports noncortical boundary definition and yields a reproducible, developmentally appropriate parcellation that generalizes across independent cohorts.

### Whole-brain connectivity reveals division-specific internal architectures

Whole-brain functional connectivity revealed an organized architecture spanning the cerebral cortex, cerebellum and subcortex, with differentiated within-division structure embedded within selective cross-division coupling (Fig. 6a). Averaged from birth to six years, both diagonal and off-diagonal blocks of the connectivity matrix were patterned rather than homogeneous, indicating that early-life functional organization was neither confined to individual divisions nor characterized by diffuse whole-brain coupling.

**Figure 6.**
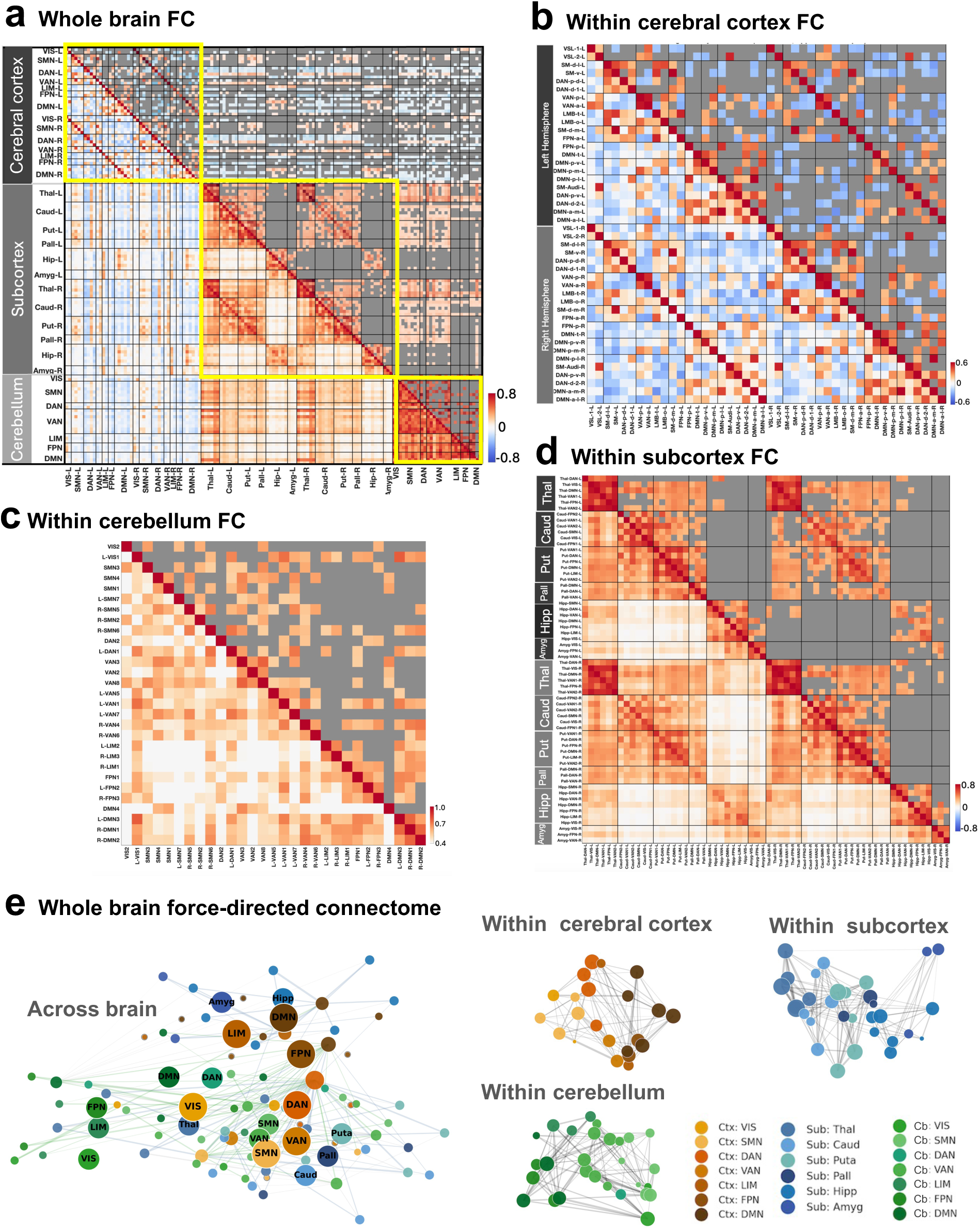
Whole-brain connectivity reveals division-specific internal architectures within a shared relational space. **a**, Whole-brain functional connectivity (FC) matrix averaged across 0-6 years, showing intrinsic interactions across the cortex, subcortex, and cerebellum. The lower triangle shows Fisher-z-transformed FC values, whereas the upper triangle shows block-wise thresholded connectivity, with values below the 60th percentile of absolute FC within each block are masked in gray to highlight the strongest retained within- and between-division couplings. Yellow boxes indicate the within-cortex, within-subcortex and within-cerebellum blocks enlarged in b-d. **b**, Within-cortex connectivity, ordered by seven large-scale functional systems and hemisphere, showing differentiated, dual-polarity organization with positive within-system and selected negative between-system relationships. **c**, Within-cerebellum connectivity, showing predominantly positive, modular interactions among functionally related territories. **d**, Within-subcortex connectivity, ordered by nucleus, showing nucleus-centered organization with stronger positive coupling within thalamic and striatal-pallidal territories and more selective hippocampal and amygdalar interactions. **e**, Force-directed embeddings constructed from the strongest retained absolute FC values. In the across-brain embedding, cortical, subcortical and cerebellar parcels occupy partially separable but overlapping connectivity neighborhoods; node size is proportional to weighted degree. Visual, somatomotor and attention-related cortical parcels are positioned near selected thalamic, striatal–pallidal and cerebellar parcels with corresponding functional affiliations, whereas hippocampal and amygdalar parcels occupy a more limbic–association-related neighborhood. Within-division embeddings show the retained connectivity structure separately for cortex, subcortex and cerebellum. Ctx, cerebral cortex; Sub, subcortex; Cb, cerebellum.

Each division exhibited a distinct internal connectivity signature. The cortical matrix showed a pronounced dual-polarity organization, with positive and negative edges comprising 47.7% and 52.3% of connections, respectively. Positive coupling was strongest within visual and somatomotor systems (mean Fisher z-transformed functional connectivity (zFC) = 0.553 and 0.402), whereas negative coupling characterized selected cross-system relationships, including visual–frontoparietal and visual-ventral-attention connections (mean zFC = −0.134 and −0.129; Fig. 6b). Cerebellar connectivity was uniformly positive in the group-average matrix, with strong within-system coupling in default-mode and ventral-attention territories (mean zFC = 0.748 and 0.720) and strong somatomotor-ventral-attention coupling (mean zFC = 0.705; Fig. 6c). The subcortex showed a nucleus-centered architecture: within-nucleus coupling was substantially stronger than between-nucleus coupling (mean zFC = 0.585 versus 0.218), with the strongest internal coupling in the thalamus and putamen (mean zFC = 0.859 and 0.626; Fig. 6d). Thus, the shared scaffold incorporated three distinct intrinsic architectures rather than a uniform connectivity pattern across divisions.

Between-division connectivity was likewise structured. Cortex–cerebellum coupling was strongest among visual- and somatomotor-related territories (mean zFC = 0.116 and 0.060), whereas cortical frontoparietal parcels showed negative coupling with several cerebellar systems. Cerebellum-subcortex coupling was predominantly positive, with the strongest relationships linking cerebellar ventral-attention and somatomotor territories to subcortical association-related parcels (Fig. 6a). A force-directed embedding of the strongest absolute FC values further showed partially separable but overlapping connectivity neighborhoods across divisions (Fig. 6e). Visual, somatomotor and attention-related cortical parcels were positioned near selected thalamic, striatal–pallidal and cerebellar parcels with corresponding affiliations, whereas hippocampal and amygdalar parcels occupied a more limbic–association-related neighborhood.

Together, these analyses show that early-life whole-brain organization is defined by selective relationships among a differentiated, dual-polarity cortical architecture, a predominantly positive cerebellar architecture and a nucleus-centered subcortical architecture.

### Development selectively refines the shared whole-brain functional scaffold

To characterize the temporal dynamics of early-life connectome reorganization, we modeled developmental trajectories of functional connectivity from birth to six years. Rather than maturing uniformly, functional connections followed heterogeneous, structure-specific developmental trajectories that were largely preserved after sex stratification, with only modest network-specific differences between females and males (Supplementary Fig. 3 and 4).

Within-division trajectories differed markedly across the three brain divisions (Extended Data Fig. 7). Cortical changes were heterogeneous in shape but generally modest in amplitude. Visual and somatomotor systems showed early declines followed by stabilization or later rebound, whereas association systems showed gradual increases, shallow nonmonotonic profiles or multiphasic changes. Subcortical trajectories were comparatively flat and less differentiated, consisting mainly of stable, mildly declining or weakly nonmonotonic profiles. Cerebellar trajectories showed larger-amplitude and more nonlinear remodeling. Somatomotor, limbic, frontoparietal and default-mode systems exhibited prominent U-shaped or rebound-like profiles, whereas visual and dorsal-attention connectivity declined more consistently. The shared scaffold was therefore refined through modest cortical change, comparatively constrained subcortical remodeling and pronounced nonlinear cerebellar reorganization.

This division-specific divergence extended to the whole-brain developmental landscape (Fig. 7a). Consistently positive coupling was most evident within the intra-subcortical and intra-cerebellar blocks, and along the within-network diagonal of intra-cortical connectivity, indicating relatively stable intra-structural cores across early life. Intra-cortical connections, however, showed the greatest diversity of sign-change patterns and trajectory profiles. Cross-structure interactions were more heterogeneous. Cortex-subcortex connectivity preserved positive coupling along several functionally corresponding axes, while selected connections shifted from negative to positive, indicating progressive strengthening along specific cortico-subcortical relationships. Cortico-cerebellar interactions showed greater reconfiguration: stable positive coupling was preserved mainly along primary cortical axes, especially visual and somatomotor systems, whereas many other pairs remained negative or shifted from positive to negative. Cerebello-subcortical interactions were more constrained and were predominantly characterized by decreasing trajectories. The connections showing the largest fitted developmental changes are summarized in Supplementary Fig. 5.

**Figure 7.**
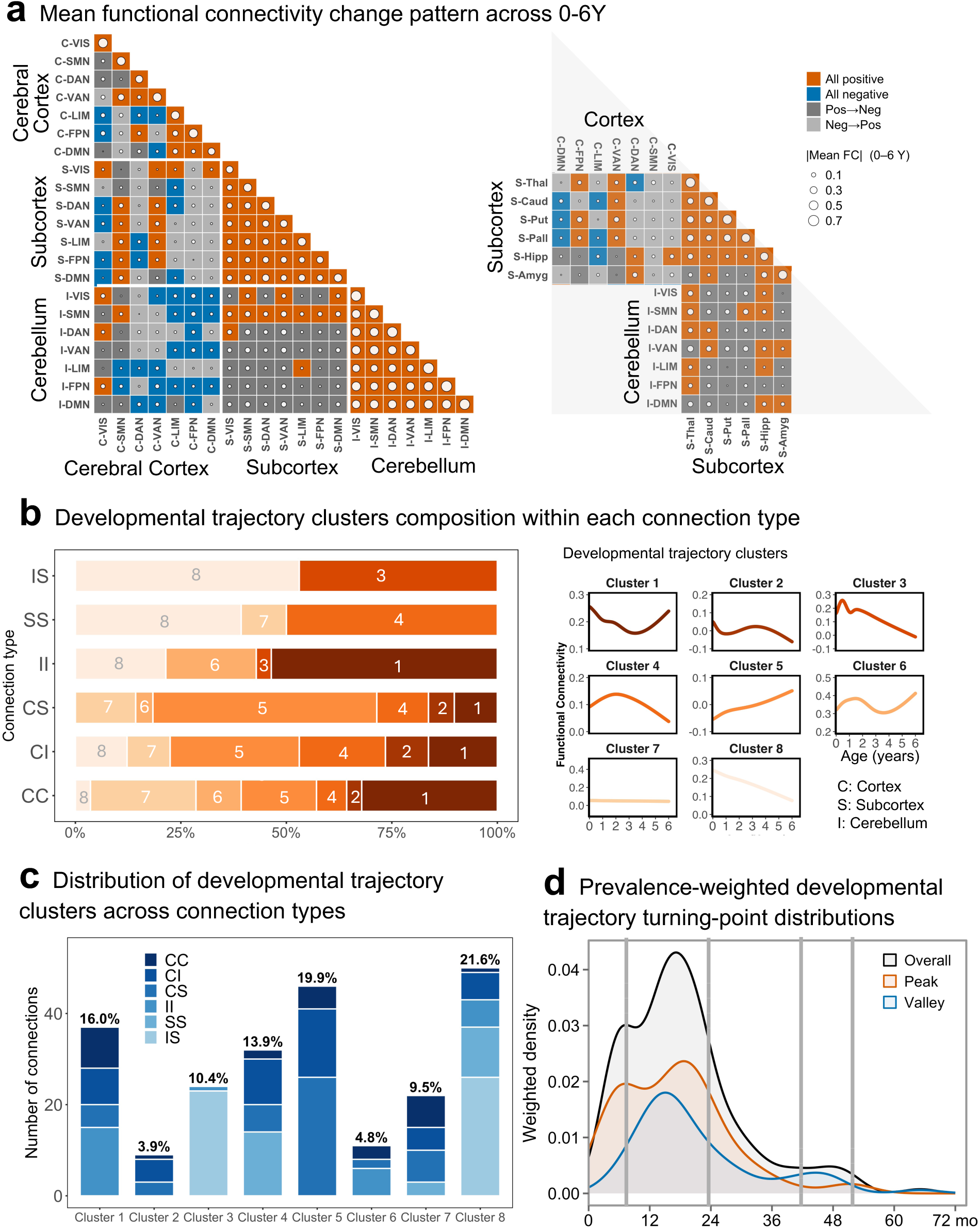
Development selectively refines the whole-brain functional scaffold through connection-specific trajectories with early-life inflections. **a,** System-level summaries of developmental functional connectivity (FC) trajectories from birth to six years. The left matrix uses a unified 21-node representation in which cerebral cortex, subcortex and cerebellum are each summarized into seven functional systems; the right matrix resolves the subcortex into individual nuclei. Tile color indicates the sign pattern of the fitted developmental trajectory across age: consistently positive, consistently negative, positive-to-negative or negative-to-positive. Circle size indicates the absolute mean Fisher z-transformed FC across the 0-6-year interval. Intra-division blocks (CC, SS, II) are dominated by stable positive coupling, whereas cross-division interfaces, particularly those involving the cerebellum, are highly dynamic and enriched for sign-switching. **b**, Distribution of eight empirically defined trajectory categories across anatomical connection classes. Stacked bars show the proportional composition of trajectory categories within each class; numbers indicate category labels. Representative GAMM-fitted shapes are shown at right. Cortex-cortex connections span nearly the full repertoire of trajectory motifs, whereas cerebellum-subcortex connections are concentrated in declining profiles. **c**, Number of connections assigned to each trajectory category, stratified by anatomical connection class. The distribution shows that trajectory patterns are not uniformly distributed across the connectome. Cortex-anchored classes occupy a broader range of trajectories, whereas cerebellum–subcortex connections show more constrained developmental profiles. **d**, Prevalence-weighted density distributions of the first detected developmental turning point across all analyzed trajectories. Peaks indicate the first positive-to-negative slope reversal, marking an early local maximum in FC trajectories, whereas valleys indicate the first negative-to-positive slope reversal, marking an early local minimum. Turning points are concentrated early in life, with an initial elevation in the first postnatal year and a dominant peak around 15-18 months. CC, cerebral cortex-cerebral cortex; CI, cerebral cortex-cerebellum; CS, cerebral cortex-subcortex; II, cerebellum-cerebellum; SS, subcortex-subcortex; IS, cerebellum-subcortex. Network abbreviations are as in Fig. 1.

Empirically defined trajectory categories recapitulated these division-specific distinctions (Fig. 7b, c). Cortex-cortex (CC) connections showed the broadest repertoire, spanning nearly all trajectory clusters and indicating the greatest diversity of intra-cerebral maturation profiles. More broadly, cortex-anchored connection classes exhibited the widest combination of sign-change types and trajectory motifs, suggesting that cortical systems provide the most differentiated axis of whole-brain FC reorganization. Cerebellum-subcortex (IS) connections were more constrained, consisting predominantly of Clusters 8 and 3, both marked by overall declining trajectories. Cerebellum-cerebellum (II) connections were enriched for Cluster 1 and other nonlinear profiles, consistent with rebound-like remodeling within the cerebellum. By contrast, cortex-subcortex (CS) and, to a lesser extent, cortex-cerebellum (CI) were enriched for Clusters 5 and 4, indicating progressive increases or shallow nonmonotonic remodeling, although CI also retained a declining component.

Finally, turning-point analysis showed that major developmental inflections were concentrated early in life (Fig. 7d). The density of first turning points rose rapidly in infancy, showed an early elevation around 6 months, and reached a broader dominant peak around 15-18 months. Thereafter, density declined sharply, with only weak later shoulders and minimal events toward the end of the 0-6-year interval. Peak-like events were more prevalent than valley-like events, although both were concentrated primarily within the first two years of life. Collectively, these results indicate that early functional maturation is organized by structure-specific trajectory profiles, with infancy and the second year of life forming a concentrated window of large-scale connectome reorganization.

### Distinct scaffold components contribute differentially to developmental and cognitive prediction

To test whether development and prospective cognition-related variation was differentially distributed across the scaffold, we trained machine-learning models to predict sex, chronological age at scanning and cognitive performance assessed 6-12 months later, using UNC-ELF seven-system functional representations of the cortex, cerebellum and subcortex (Fig. 8).

**Figure 8.**
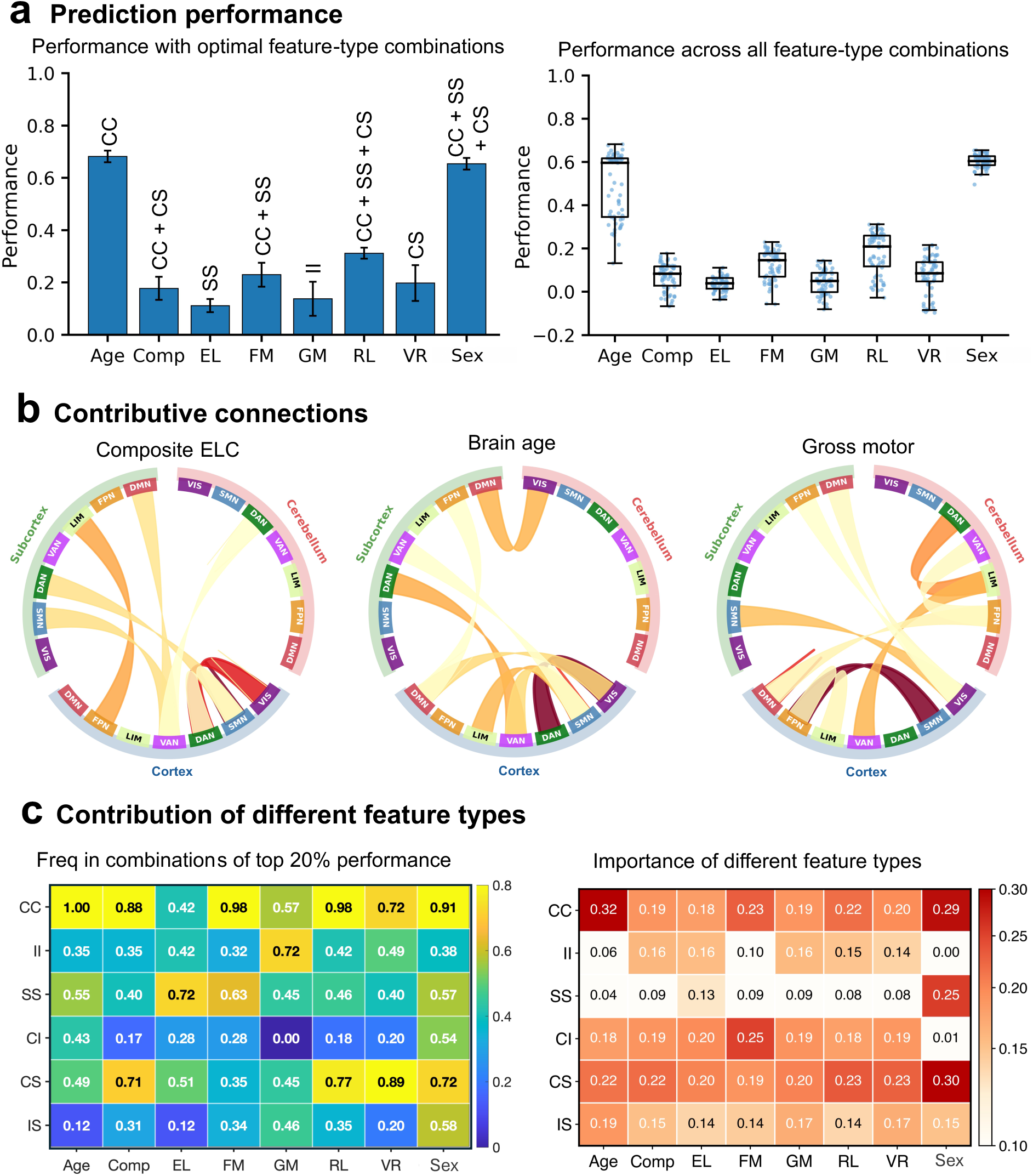
Distinct scaffold components differentially encode developmental and cognitive variation. **a**, Prediction performance for age at scan, sex and prospective cognitive outcomes assessed 6-12 months after scanning. Left, bar plots show mean performance achieved by the best-performing combination of functional connectivity (FC) classes for each target, with the selected FC-class combination annotated above each bar. Right, box plots show prediction performance across all 63 non-empty combinations of the six FC classes. Performance is reported as Pearson correlation coefficient for age and continuous cognitive outcomes, and area under the receiver operating characteristic curve for sex classification. Error bars indicate s.d. across repeated cross-validation folds. **b**, Contributive FC connections for representative prediction targets, including Early Learning Composite, brain age and gross motor performance. Connections are mapped according to structural assignment and seven-network identity. Ribbon thickness and opacity indicate relative feature importance, showing that different targets rely on distinct whole-brain connectivity patterns. **c**, Target-specific contribution of FC classes. Left, heatmap showing how frequently each FC class appears among the top 20% performing feature-class combinations for each target. Right, heatmap showing normalized summed feature importance for each FC class within each target. Cortico-cortical and cortico-subcortical features contribute broadly across targets, whereas selected domains show more specific contributions, including cerebellum-cerebellum features for gross motor prediction and subcortex-subcortex features for expressive language and sex classification. For box plots, center line, median; box limits, upper and lower quartiles; whiskers, 1.5× interquartile range; points, individual FC-class combinations. CC, cortex-cortex; II, cerebellum-cerebellum; SS, subcortex-subcortex; CI, cortex-cerebellum; CS, cortex-subcortex; IS, cerebellum-subcortex. Comp, Early Learning Composite; EL, expressive language; FM, fine motor; GM, gross motor; RL, receptive language; VR, visual reception.

Prediction was strongest for chronological age, which was captured most accurately by cortex-cortex connectivity (CC) alone (Pearson *r* = 0.68; Fig. 8a, b), indicating that intra-cortical network organization carries a substantial component of early maturational variation. Sex was also classified above chance, with the highest performance achieved using a broader model combining cortex-cortex, subcortex-subcortex and cortex-subcortex features (CC + SS + CS; AUC = 0.66), suggesting that early sex-related differences are distributed across cerebral, subcortical and cortico-subcortical organization rather than confined to a single connection class.

Prospective cognitive prediction was more modest but varied across domains. Receptive language showed the strongest prediction among cognitive measures (*r* = 0.31; CC + SS + CS), followed by fine motor performance (*r* = 0.23; CC + SS), visual reception (*r* = 0.20; CS) and the Early Learning Composite (*r* = 0.18; CC + CS). Gross motor performance was best predicted by intra-cerebellar features (II; *r* = 0.14), whereas expressive language showed the weakest positive prediction (SS; *r* = 0.10). Across all 63 FC-class combinations, age and sex showed the strongest overall prediction performance, while receptive language was the most consistently predicted cognitive domain (Fig. 8b). Compared with an adult-derived whole-brain reference scheme, the Early Life Whole-Brain Functional Parcellation also yielded more favorable prediction performance across multiple developmental and behavioral targets (Extended Data Fig. 9).

Feature-importance analyses further showed that predictive information was distributed across the whole-brain scaffold in a target-specific manner (Fig. 8c, d). At the macroscale, CC features showed the broadest involvement across targets and dominated age prediction, whereas CS features contributed prominently to prediction of the Early Learning Composite, receptive language, visual reception and sex. By contrast, SS features were more selectively enriched for expressive language and fine motor prediction, and II features were most prominent for gross motor prediction. Connectogram visualizations of the most informative individual features revealed distinct connection patterns across targets (Fig. 8c). Age prediction was dominated by intra-cerebral links involving visual, somatomotor and attention systems, suggesting that maturational variation is strongly represented in coordinated sensory-attention architecture. Prediction of the Early Learning Composite relied on a broader integrative pattern, combining focal intra-cerebral connections with distributed cortico-subcortical links centered on visual, somatomotor and attention networks. Gross motor prediction, by contrast, showed a more cerebellar-anchored pattern, with prominent intra-cerebellar and cerebello-cortical links converging on somatomotor and attention systems.

These predictive patterns indicate that the early-life whole-brain scaffold carries target-specific developmental and cognitive information. Cortical and cortico-subcortical connections carry the broadest predictive signal, whereas subcortical and cerebellar components contribute more selectively to motor and specific cognitive domains.

## DISCUSSION

Early human functional development has often been interpreted either through cortical network maturation or through analyses of individual noncortical structures. Our findings support a more integrated view: early neurodevelopment is already organized at the whole-brain level. The observation that all seven cortical functional systems are represented beyond the cerebral cortex, together with explicit correspondences across cortex, cerebellum and subcortex, broadens the traditional view of a mature-like cortex surrounded by loosely connected auxiliary structures. Instead, early functional organization appears to be anchored by a shared scaffold spanning the major brain divisions, while preserving division-specific patterns.

Existing functional parcellations have been highly informative, but they have generally been constructed through fragmented routes, either by extending cortical schemes to noncortical structures or by parcellating cerebellum and subcortex independently^6,8,15,16^. In early life, this fragmentation is especially consequential because parcellation choice shapes measured connectivity and downstream inference, making studies across separately derived schemes less straightforward^20^. This issue is amplified by infant-specific evidence showing that age-appropriate parcellations are needed to capture coordinated developmental changes^3,10^. By deriving cortical, cerebellar and subcortical parcellations from common cohorts using a shared analytical strategy with structure-adapted connectivity features, our framework places local organization and cross-structure correspondence within the same developmental coordinate system. The reproducibility and benchmarking results further indicate that this unified UNC-ELF parcellation is stable and developmentally appropriate, supporting the conclusion that early whole-brain functional organization is both mappable and age-specialized^2,10,11,21^.

Within this unified framework, early whole-brain functional organization across the first six years of life emerges as a shared but nonuniform multi-structure scaffold, with each division contributing a distinct mode of integration. The cerebral cortex shows the most differentiated and dual-polarity architecture, consistent with canonical network segregation and a developing primary-to-transmodal hierarchy^6,21,22^. The cerebellum expresses the scaffold in a graded and predominantly positive form, with admixture concentrated along primary-to-higher-order axes. This pattern suggests that the cerebellum represents cerebral functional systems through graded topographic representation rather than sharply segregated one-to-one modules, in line with motor-to-transmodal cerebellar gradients and shared computational principles across motor and non-motor domains^8,23–26^. It also fits developmental evidence that primary cerebello-cortical coupling is established early, whereas higher-order cerebellar participation continues to refine across infancy and early childhood. The subcortex follows a different logic: an anatomically constrained yet highly integrative mosaic in which admixture is nucleus-specific and likely suggests integrative territories with distinct circuit roles^16,27^. Thalamic and pallidal territories preferentially combine sensory and association correspondence, striatal territories show stronger attention-limbic-sensorimotor overlap, and hippocampal and amygdalar parcels bridge primary/attention and limbic/association systems^4,12,28,29^. Together, these findings suggest complementary modes of integration: hierarchical differentiation in cortex, graded coordination in cerebellum, and nucleus-constrained routing in subcortex.

The FC architecture reinforces this interpretation. The whole-brain matrix and force-directed embedding revealed an ordered relational space rather than a diffuse mixture of regions. Sensorimotor- and attention-related cortical systems were positioned near thalamic and striatal-pallidal parcels, whereas hippocampal and amygdalar parcels occupied a more limbic-association periphery. This organization suggests that cerebellar and subcortical territories are not simply peripheral extension of cortical functional systems, but integral components of early whole-brain architecture that participate in central perception- and action-related systems while preserving more specialized territories for affective and higher-order association functions.

From a developmental perspective, our findings suggest that early maturation proceeds less by constructing a new architecture than by coordinately refining an established scaffold. Although recognizable organization is present near birth, different systems mature on distinct timescales^1,30–32^. Within this framework, intra-structural cores appear comparatively preserved, whereas cross-structure interfaces remain more developmentally plastic. This preservation is not uniform: cortex-cortex and cortex-anchored connections show the broadest range of sign patterns and trajectory profiles, indicating that cortical systems provide the most differentiated axis of whole-brain FC reorganization, consistent with interactive specialization accounts in which development reflects progressive reweighting among already differentiated systems^33^. Major cross-structure interfaces, however, follow distinct developmental rules. Cortico-subcortical coupling appears comparatively aligned and progressively coordinated, consistent with evidence that thalamocortical organization is already structured early in life^3,4,34,35^. By contrast, cerebellar interfaces emerge as a prominent locus of selective recalibration: stable positive cortico-cerebellar coupling is concentrated mainly along visual and somatomotor axes, whereas many non-primary cerebello-cortical, and especially cerebello-subcortical, relationships show declining or sign-switching trajectories. This pattern argues against a simple delay model and instead suggests progressive differentiation of cerebellar participation in whole-brain circuitry^5,36^. Notably, major FC inflections were concentrated within infancy and the second year of life, indicating that large-scale connectome remodeling is strongly front-loaded across 0-6 years. Together, these findings point to an early, temporally concentrated window in which a relatively preserved internal scaffold is selectively refined through reconfiguration of cross-structure communication.

Our predictive analyses further suggest that developmental phenotypes are distributed across different components of this scaffold rather than captured by a single global FC factor. Age-related information was concentrated primarily in within-cerebral circuitry, implying that maturational variation is strongly encoded in refinement of cortical architecture^1,32^. By contrast, prospective cognitive outcomes were not anchored to cortex alone: cortico-subcortical features repeatedly emerged among the most informative predictors, suggesting that later cognition variation may be partly reflected in how cortical systems are integrated with subcortical circuitry during early life^4,29,34^. Gross motor prediction showed the clearest prominence of within-cerebellar organization, reinforcing the view that the cerebellum is already an important component of early sensorimotor developmental architecture^5,36^. More broadly, these findings suggest that early neurodevelopmental variation is encoded through a structured allocation of information across cortical, subcortical and cerebellar interactions.

Several limitations define the scope of these findings. The present framework is based on resting-state functional connectivity and therefore characterizes coordinated organization rather than causal or anatomical pathways. In addition, prospective cognitive prediction effects were modest and were evaluated through internal validation rather than external replication, warranting appropriate caution^37^. These limitations clarify the next step: embedding this unified functional scaffold within a multimodal developmental framework incorporating structural connectivity, morphology and myeloarchitectural change^38^. Integrating these biological features could clarify how the shared functional scaffold and its target-specific remodeling arise from underlying maturation. Future work should also extend validation to atypical or high-risk cohorts to determine whether whole-brain parcellation provides added value over structure-specific parcellations for detecting developmentally relevant alterations.

## MATERIALS AND METHODS

### Participants and imaging acquisition

This study included 372 participants contributing 913 MRI scans from two longitudinal infant and early childhood cohorts: the Multi-visit Advanced Pediatric Brain Imaging Study (MAP)^39^ and the UNC/UMN Baby Connectome Project (BCP)^40^. The experimental protocols were approved by the Institutional Review Board of the University of North Carolina at Chapel Hill, and written informed consent was obtained from a legal guardian for each participant. Participants were recruited through existing registries at the University of North Carolina (UNC) and the University of Minnesota (UMN), including statewide birth records, as well as broader community resources such as community centers and targeted daycare programs, with the goal of approximating the racial, ethnic, and socioeconomic diversity of the US population.

The MAP cohort included 76 participants with 316 longitudinal scans acquired from birth to 6 years of age on 3T Siemens scanners. T1-weighted images were acquired with TR = 1,820 *ms*, TE = 4.38 *ms*, and resolution = 1 × 1 × 1 *mm*^3^. Resting-state fMRI (rs-fMRI) data were acquired using the EPI sequence with TR = 2,000 *ms*, TE = 32 *ms*, resolution = 4 × 4 × 4 *mm*^3^, and a total of 150 volumes. To maintain image quality, infants younger than 2 years were scanned during natural sleep, while older children were imaged while awake and watching movies. The BCP cohort included 296 participants with 597 longitudinal scans acquired between birth and 6 years of age, also on 3T Siemens MRI scanners. T1-weighted images were acquired with TR = 2,400 *ms*, TE = 2.24 *ms*, and resolution = 0.8 × 0.8 × 0.8 *mm*^3^. Rs-fMRI data were collected using an EPI sequence with TR = 800 *ms*, TE = 37 *ms*, resolution = 2 × 2 × 2 *mm*^3^, and 420 volumes. Children younger than 3 years were scanned during natural sleep, whereas older children were scanned while awake and watching movies. In addition, 21 BCP participants aged 3–6 years underwent both awake and sleep rs-fMRI acquisitions, enabling within-cohort modeling of scan-state translation.

A subset of the BCP cohort also contributed behavioral data. Specifically, 364 scans from 212 participants aged 3-61 months were linked to prospective assessments with the Mullen Scales of Early Learning (MSEL)^41^, administered 6-12 months after imaging, yielding the Early Learning Composite and domain-specific scores for gross motor, fine motor, expressive language, receptive language and visual reception. In this study, the final Early Life Whole-Brain Functional Parcellation was derived using the combined BCP and MAP datasets to improve age coverage and create a more balanced sample across developmental stages, while also increasing overall sample size and the robustness of parcel estimation. Subsequent developmental, connectivity, and prediction analyses were performed using the BCP dataset only, because BCP provides the largest harmonized longitudinal sample with consistent imaging, demographic, and behavioral measures, enabling unbiased evaluation of age-related trajectories and brain-behavior associations within a single well-characterized cohort.

External validation used three public cohorts, Pixar^17^ (42-72 months), Calgary^18^ (31-83 months) and dHCP^19^ (39-45 weeks postmenstrual age, restricted to scans acquired around birth), chosen to test whether the Early Life Whole-Brain Functional Parcellation generalizes across acquisition sites and imaging conditions. Importantly, the parcellation was applied directly to each external dataset without harmonization, providing a stringent test of its out-of-sample generalizability and robustness across acquisition contexts. Pixar participants were recruited from the local community in Massachusetts, USA, and Calgary participants were recruited from the local community in Calgary, Canada. The dHCP sample was obtained from the Developing Human Connectome Project, with MRI acquired during natural sleep. Pixar and Calgary data were acquired under awake viewing conditions, including movie watching in Pixar and passive video viewing in Calgary. Demographic characteristics and imaging parameters for these external datasets are summarized in Supplementary Table 1 and Supplementary Table 2 and shown in Extended Data Fig. 1.

### MRI preprocessing and structure-specific signal extraction

#### Cortical surface-based preprocessing

For each scan, structural MRI was processed with the infant Brain Extraction and Analysis Toolbox (iBEAT v2.0)^42^. Processing included linear alignment of T2-weighted images to the corresponding T1-weighted images, bias-field correction, infant-specific learning-based skull stripping, cerebellum extraction, and tissue segmentation into gray matter, white matter, and cerebrospinal fluid (CSF). Infant-dedicated cortical surface reconstruction was then performed to generate the inner surface at the gray-white matter boundary, the pial surface at the gray matter-CSF boundary, and a middle cortical surface positioned within the cortical gray matter and equidistant from the inner and outer surfaces at corresponding vertices^43^. Resting-state fMRI data were preprocessed using FSL^44^, including removal of the first 10 volumes, motion correction, and temporal band-pass filtering (0.01–0.08 Hz). Functional images were rigidly aligned to the T1-weighted images using boundary-based registration^45^. Nuisance signals, including mean white matter signal, mean CSF signal, and 24 motion parameters, were regressed out. The cleaned rs-fMRI data were then projected onto the reconstructed middle cortical surface, registered and resampled to the FreeSurfer fsaverage5 surface^46^ with 10,242 vertices on each hemisphere, and smoothed with a 2-mm full-width at half-maximum kernel in both volume and surface domains using Freesurfer and Connectome Workbench, following prior recommendations^47^. Scans with excessive motion, defined as mean framewise displacement > 0.5 mm, were excluded from all analyses.

#### Cerebellar and subcortical fMRI preprocessing

For cerebellar analyses, the cerebellar mask derived from iBEAT-based structural preprocessing was used to extract the cerebellar T1-weighted image, which was then registered with ANTs^48^ to the 24-month template in the longitudinally consistent 4D infant cerebellum atlas^49^. We selected the 24-month template as an age-appropriate reference because it provides a developmentally intermediate common space within the 0-6-year window, with more stable cerebellar anatomy and tissue contrast than young-infant templates while avoiding bias toward older-child morphology. The resulting deformation fields were applied to the corresponding preprocessed rs-fMRI data, placing all individual cerebella into a common anatomical space. This age-independent framework ensured a consistent number and spatial arrangement of gray matter voxels across participants, enabling construction of a single comparable cerebellar parcellation across the full age range. A cerebellar gray matter mask derived from the 24-month atlas was then used to extract voxel-wise fMRI signals for downstream parcellation.

Subcortical rs-fMRI data were processed in parallel using the same age-independent strategy. Each individual T1-weighted image and corresponding preprocessed rs-fMRI scan was registered with ANTs to the 24-month template of the longitudinally consistent UNC 4D infant brain atlas^50^. Six bilateral subcortical structures were extracted on the basis of the UNC infant atlas parcellation: thalamus, caudate, putamen, pallidum, hippocampus, and amygdala. Functional signals within these structures were used for downstream subcortical parcellation and whole-brain analyses.

### Functional feature construction for whole-brain parcellation

#### Age-independent feature construction

For each scan, Pearson correlation-based functional connectivity (FC) matrices were computed for all feature types used in whole-brain parcellation, including intra-cerebral, intra-cerebellar, intra-subcortical, cerebello-cerebral, between-subcortical, and subcortical-cerebral FC. Each scan-level FC matrix was binarized by retaining the strongest 10% of connections, and these binarized matrices were then aggregated across all scans to generate age-independent group-level FC matrices. For each group-level FC matrix, Laplacian maps were computed using the BrainSpace toolbox^51^ with the normalized-angle (‘na’) kernel and used as clustering features for parcellation.

For cerebral cortex parcellation, Laplacian-map features were derived only from intra-cerebral FC, because cortico-cortical connectivity provides the clearest representation of cortical functional systems while preserving the cerebral cortex as an independent anchor for subsequent cross-structure correspondence analyses. For cerebellum parcellation, both intra-cerebellar FC and cerebello-cerebral FC were included, allowing clustering to capture both intrinsic cerebellar organization and its functional coupling with the cerebral cortex. For each subcortical structure, including the thalamus, caudate, putamen, pallidum, hippocampus, and amygdala, three FC types were incorporated: intra-subcortical FC within the target structure, subcortical-subcortical FC between the target structure and the remaining subcortical structures, and subcortical-cerebral FC between the target structure and the cortical surface. By combining intrinsic and cross-structure FC features, cerebellar and subcortical parcellations capture local organization together with their broader circuit embedding, producing maps that are both regionally specific and whole-brain informed. After Laplacian-map computation, each retained dimension was z-score normalized, and the resulting maps were concatenated within each structure to form the final clustering feature set. Accordingly, cerebellar features consisted of concatenated intra-cerebellar and cerebello-cerebral Laplacian maps, whereas subcortical features consisted of concatenated intra-subcortical, between-subcortical, and subcortical-cerebral Laplacian maps.

#### Age-dependent weighted feature construction

Age-dependent parcellation features were generated using the same FC definitions, top-10% binarization, Laplacian mapping, and feature concatenation procedures as in the age-independent analysis. The only difference was that group-level FC matrices were estimated using age-weighted aggregation rather than simple averaging across all scans, thereby improving longitudinal consistency while accommodating inter-individual variation. Early life was divided into nine overlapping age groups: 0M, 3M, 6M, 9M, 12M, 18M, 24M, 36M, and 72M, M for months. For each age group, the contribution of the *i*th scan’s binarized FC matrix *FC_i_* to this age group was determined by a trapezoid membership function,

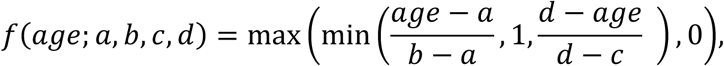

where *a*, *b*, *c*, *d* define the support of the target age bin. For degenerate cases in which *a* = *b* or *c* = *d*, the corresponding ramp denominator was not evaluated and treated as a vertical boundary: when *a* = *b*, membership was set to 1 in [*a*, *b*]; when *c* = *d*, membership was set to 1 in [*c*, *d*]. The age-bin parameters were set as follows (in days): 0M-[0, 0, 60, 90], 3M-[60, 60, 120, 150], 6M-[120, 150, 210, 240], 9M-[210, 240, 300, 330], 12M-[300, 330, 420, 450], 18M-[420, 465, 600, 660], 24M-[600, 660, 780, 840], 36M-[750, 840, 1,080, 1,080], 72M-[1,080, 1,200, 2,250, 2,250]. For a given age group, membership values were normalized across all scans with positive membership,

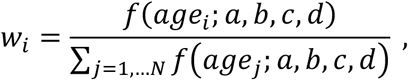

where *N* denotes the number of scans whose ages fell within the corresponding age bin. The corresponding age-specific group-level FC matrix was then computed as *FC* = ∑*_i_*_=1,…*N*_ *w_i_ FC_i_*. This weighted aggregation was applied to all FC types used for Laplacian map computation, yielding age-specific group-level clustering features.

### Harmonization of clustering features

#### Awake-to-sleep harmonization

A major source of heterogeneity between datasets was scan state: in BCP, children younger than 3 years were predominantly scanned during natural sleep, whereas older children were scanned during wakefulness; in MAP, the corresponding transition occurred at 2 years. To reduce this state-related discrepancy, we developed a learning-based harmonization procedure to transform awake Laplacian features into a sleep-like representation.

The harmonization model was trained using 21 BCP participants who underwent paired awake and sleep rs-fMRI acquisitions. To enhance predictive performance and improve model generalizability, we implemented data augmentation by partitioning each rs-fMRI scan into three non-overlapping time windows and adding random zero-mean Gaussian noise to the functional connectivity matrices computed from these windows. Each augmented segment was treated as an additional observation from the same participant and included in the training dataset. For each awake-sleep scan pair, Laplacian maps *F*^awake^ and *F*^sleep^ were computed with dimension of *V* × 50, where *V* denotes the number of cortical surface vertices or cerebellar/subcortical voxels, depending on the feature set under consideration. To predict each element of *F*^sleep^, support vector regression^52^ (SVR) was trained with the corresponding awake Laplacian map as input. For element (*v*, *j*) in *F*^sleep^, the input feature was *v*^th^ row of *F*^awake^. Model hyperparameters were selected by leave-one-out cross-validation on the paired dataset.

For clustering feature preparation, state-specific Laplacian maps were generated within each dataset from group-level FC matrices. The sleep-state feature *F*^1^ was derived from FC averaged across age groups acquired predominantly during sleep: 0M-36M in BCP and 0M-24M in MAP. Awake-state features *F*^2^ were derived from age groups acquired predominantly during wakefulness: the 72M group in BCP, and the 36M and 72M groups separately in MAP. The trained SVR model was then applied to each awake-state feature to predict its sleep-state counterpart. Thus, the BCP 72M feature was transformed into 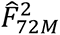, whereas the MAP 36M and 72M features were transformed separately into 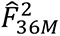 and 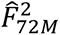. The final age-independent clustering feature was computed as an age-group-balanced weighted average, with weights reflecting the proportion of age groups represented by each state-specific feature. Because each dataset included nine age groups, the sleep-state and harmonized awake-state features were weighted as (8/9) and (1/9) in BCP, and as (7/9), (1/9), and (1/9) in MAP, respectively:

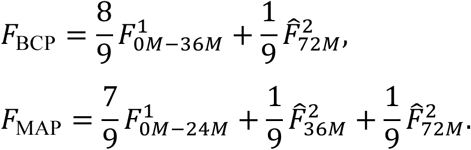

This procedure harmonized awake-derived features into the sleep-state feature space and increased cross-dataset agreement of Laplacian embedding features across all major feature types (Supplementary Fig. 1).

#### Site-effect assessment

We further evaluated the potential contribution of site-related effects to the clustering features. Matched Laplacian-derived features from BCP and MAP showed high cross-dataset similarity, with Pearson correlations generally exceeding 0.90 for corresponding features (Supplementary Fig. 1). Given this strong concordance, together with the improvement after awake-to-sleep harmonization, we did not apply an additional site-harmonization procedure.

### Cerebral cortex, cerebellum, and subcortex clustering

For each set of Laplacian-map features, whether derived from the age-independent or age-dependent framework, parcellation was performed using k-means clustering^53^ in MATLAB R2022b. Clustering was run with a maximum of 1,000 iterations and 200 replicates, and the solution with the smallest within-cluster sum of distances to cluster centroids was retained. BCP and MAP were each partitioned into test-retest subsets with comparable age-group and sex composition. The BCP and MAP test subsets were then merged to generate the full test cohort, and the corresponding retest subsets were merged to generate the full retest cohort. For each brain division, clustering was optimized by tuning the number of retained Laplacian feature dimensions on the basis of test-retest reproducibility between the split-half test and retest cohorts and the silhouette index^54^.

Cortical parcellations were visualized on the cortical surface template. Cerebellar parcellations were visualized on the 24-month infant cerebellum template, and were additionally projected onto the SUIT cerebellar flat map^55,56^. For SUIT visualization, the infant cerebellum template was first registered to the SUIT template using ANTs, and the resulting deformation fields were applied to transform parcellation maps from native cerebellar space into SUIT space. Subcortical parcellations were visualized on reconstructed surface meshes for each subcortical structure. These meshes were generated from binary masks by extracting three-dimensional iso-surfaces. A detailed schematic of the full whole-brain parcellation workflow is provided in Extended Data Fig. 1.

### Benchmarking against existing parcellations

#### Parcellation matching and Dice correspondence analysis

To evaluate the developmental appropriateness of the Early Life Whole-Brain Functional Parcellation, we benchmarked our age-independent functional parcellation against widely used existing functional parcellations at matched or approximately matched spatial scales across the cerebral cortex, cerebellum and subcortex. In the cerebral cortex, the 22-network Early Life Whole-Brain Functional Parcellation was collapsed to 17 and 7 networks and compared with the adult Yeo-17 and Yeo-7 functional parcellations^6^, respectively. In the cerebellum, the Early Life 54-parcel, 29-parcel and 54-to-7 functional cerebellar parcellations were compared with the Nettekoven-68, Nettekoven-32, Buckner-17 and Buckner-7 functional parcellations^8,15^. To enable resolution-matched comparisons, the Nettekoven 68- and 32-parcel functional parcellations were merged into the Early Life 54- and 29-parcel label spaces using a Dice-overlap-based assignment procedure. Specifically, a Dice similarity matrix^57^ was computed between all source and target parcels, then each source parcel was assigned to the target parcel with the greatest Dice overlap. When necessary, a repair step was applied to reassign parcels from overrepresented targets to previously unassigned targets while minimizing loss of Dice similarity. Source voxels were then relabeled according to the final assignments to generate the merged cerebellar functional parcellations.

For the subcortex, the Early Life functional parcellation was compared with Tian^16^ Scale IV both in its native form and after harmonization to a matched 46-label representation, defined within each anatomical structure by the minimum number of subdivisions shared by the two parcellations. This matched representation comprised 6 thalamic, 4 caudate, 4 putamen, 2 pallidal, 5 hippocampal and 2 amygdalar labels per hemisphere. For paired bilateral visualization, homologous right-hemisphere labels were relabeled to the corresponding left-hemisphere identities.

To quantify spatial correspondence with established adult reference systems, we additionally computed per-network Dice overlap between Early Life and adult functional parcellations after collapsing them to comparable network definitions. In the cerebral cortex, Dice overlap was computed between the Early Life 22-to-17 mapping and Yeo-17 separately in the left and right hemispheres on the fsaverage5 surface and then averaged across hemispheres for each network. Networks were further grouped into the seven canonical large-scale systems, and system-level trends were summarized by averaging Dice values within each system and testing their monotonic variation across the cortical hierarchy using Spearman rank correlation. In the cerebellum, Dice overlap was computed between the Early Life 54-to-7 functional parcellation and Buckner-7 in volumetric space using matched network identities (VIS, SMN, DAN, VAN, LIM, FPN and DMN). These analyses were used to quantify the correspondence between early life brain organization and established adult canonical systems across functional hierarchies.

#### Functional homogeneity analysis

Functional homogeneity was computed using the same procedure for the cerebral cortex, cerebellum and subcortex^7^. For each subject and each functional parcellation, we extracted the fMRI time series of all grayordinates within each parcel, where grayordinates corresponded to surface vertices in the cerebral cortex and voxels in the cerebellum and subcortex. For parcel *p*, we calculated the parcel-mean time series, *x̄_p_*, correlated each grayordinate time series, *x_v_*, with *x̄_p_*, applied Fisher’s *z*-transformation, and averaged the resulting values within the parcel. Averaging was performed in Fisher-z space because Fisher transformation reduces bias from the bounded scale of correlation coefficients and provides a variance-stabilized scale for aggregation. Whole-structure homogeneity was then defined as the parcel-size-weighted mean of parcel homogeneity values in Fisher-*z* space, followed by inverse transformation back to correlation space. This yielded one homogeneity value per subject for each functional parcellation. For the subcortex, homogeneity was additionally summarized within each anatomical structure.

Because each subject was evaluated under every functional parcellation, statistical comparisons were performed within subject. One-sided paired *t*-tests were used to test whether the Early Life Whole-Brain Functional Parcellation showed greater homogeneity than the corresponding matched existing functional parcellation in each comparison. In the subcortex, comparisons were performed for both the native and matched representations, including structure-wise analyses within the thalamus, caudate, putamen, pallidum, hippocampus and amygdala. All *p* values reported in Fig. 5 are one-sided.

#### Cross-structure boundary contrast analysis

To test whether integrated parcellations better captured cross-structure functional boundaries than parcellations derived from within-structure FC alone, we computed boundary contrast using cross-structure FC profiles. For each subject, cerebellar or subcortical voxels were represented by their FC profiles to cortical parcels defined using either the 22- or 36-network cortical parcellation. Boundary contrast was defined as the mean similarity between voxels within the same parcel minus the mean similarity between voxels lying across parcel boundaries. Higher values therefore indicate sharper separation of cross-structure FC profiles at parcel borders. Boundary contrast was computed for integrated UNC-ELF parcellations and resolution-matched parcellations derived from within-structure-FC only. Parcellations were compared within subject using paired tests, with FDR correction applied across matched comparisons.

#### Morphology- and tractography-based homogeneity

To further benchmark cortical functional parcellations using independent structural criteria, we evaluated morphology-based homogeneity and tractography-based homogeneity across parcellation schemes. These analyses were restricted to cerebral cortical parcellations because the selected structural benchmarks are best established for the cortical sheet. Surface-based morphological features, including sulcal depth, cortical thickness, surface area and curvature, provide spatially resolved measures of cortical structural coherence, but comparable descriptors are not directly defined for cerebellar and subcortical parcels within the same framework. Likewise, tractography-derived connectivity fingerprints are informative for cortical parcels but are less reliable for small subcortical nuclei and cerebellar parcels. Thus, morphology- and tractography-based homogeneity was used as an independent structural validation of cortical parcellation quality, complementary to functional homogeneity analyses conducted across all brain divisions.

For morphology-based homogeneity, vertex-wise cortical morphological features, including sulcal depth, cortical thickness, surface area and curvature, were first z-scored across vertices within each subject. For each parcel, the variance of each feature across vertices was computed and then averaged across features to obtain a parcel-level morphology variance. Cerebral-cortex morphology-based homogeneity was summarized as the parcel-size-weighted mean of these parcel-level variance values across all parcels, yielding one value per subject for each parcellation. Lower values indicate greater within-parcel morphological homogeneity.

For tractography-based homogeneity, we used preprocessed BCP fiber tractography data obtained directly from the Fiber Data Hub^58^ (https://brain.labsolver.org/mds/data-hcp/lifespan.html). Tractography-derived connectivity fingerprints were obtained for sampled voxels or vertices within each cortical parcel, and pairwise similarity of these fingerprints was computed within each parcel. Parcel-level tractography homogeneity was defined as the mean within-parcel similarity and then summarized at the whole-cortex level as the parcel-size-weighted mean across parcels, yielding one value per subject for each parcellation. Higher values indicate greater within-parcel tractographic homogeneity.

These analyses were used to compare the principal cortical resolution of the Early Life cerebral cortex functional parcellation (22-network) and its derived 22-to-17 and 22-to-7 mappings against the adult Yeo-17 and Yeo-7 functional parcellations. Statistical comparisons were performed using one-sided paired *t*-tests.

### Assignment of cerebellar and subcortical networks to cortical network and functional families

#### Cross-structure assignment

To characterize functional correspondence among the cerebral cortex, cerebellum and subcortex, we used a cortex-centered, bootstrap-based assignment procedure. The goal was to label each cerebellar and subcortical network according to the cortical system with which it showed the most reproducible and selective functional correspondence. The procedure included five main steps: building cross-structure FC profiles, collapsing cortical families into seven systems, applying double z-score normalization, assigning network labels, and summarizing bootstrap statistics. To establish the functional reference framework, we defined a bilateral cortical scaffold comprising seven large-scale cortical systems. Functional parcellation was performed separately in the left and right cerebral hemispheres, yielding 22 cortical parcels per hemisphere. Homologous left- and right-hemisphere parcels were matched according to interhemispheric symmetry to form 22 bilateral cortical functional families. These families were further grouped into seven large-scale cortical systems according to their surface topography and spatial correspondence with the Yeo 7-network organization. This bilateral seven-system scaffold served as the reference for assigning cerebellar and subcortical networks. Of note, this seven-system scaffold was therefore not an application of the adult Yeo-7 atlas to early-life data, but a coarse-grained representation of the Early Life 22-network cortical solution. The Yeo 7-network organization was used only as an external canonical reference for naming and aligning large-scale systems. The developmental specificity of the resulting UNC-ELF parcellation was evaluated separately through spatial-overlap and functional-homogeneity comparisons with adult Yeo-7 and Yeo-17 reference maps.

#### Step 1: Build group-level cross-structure FC profiles

In each bootstrap iteration, scans were resampled in a balanced manner with respect to age group and sex. For each selected scan, we used the FC matrix between cerebellar or subcortical networks and cortical parcels. The scan-level FC matrix was thresholded at the scan-specific 90th percentile and binarized. This step retained each scan’s strongest cross-structure connections while reducing the influence of individual differences in absolute FC magnitude. The binarized FC matrices were then averaged across resampled scans to generate a group-level cross-structure FC profile. Thus, each value in the group-level matrix represented the proportion of scans showing a strong connection between a given cerebellar or subcortical network and a given cortical parcel.

#### Step 2: Collapse cortical families into seven cortical systems

The group-level FC matrix initially described connectivity from each cerebellar or subcortical network to 22 cortical families in the left hemisphere and 22 cortical families in the right hemisphere. Within each hemisphere, the 22 cortical families were collapsed into seven cortical systems by averaging families belonging to the same system. This produced one seven-system correspondence profile for the left hemisphere and one for the right hemisphere. These two hemispheric profiles were then combined using softmax pooling:

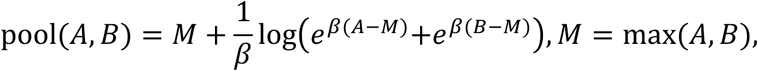

where *A* and *B* denote the left- and right-hemisphere correspondence profiles, respectively, and *β* = 1.5. The maximum *M* is subtracted for numerical stability. This operation provides a smooth max-like integration of the two hemispheres, balancing hemispheric specificity with bilateral support. It accommodates asymmetric cross-structure correspondence by preserving strong unilateral signals that would be diluted by simple averaging, while avoiding the noise sensitivity of a hard maximum.

#### Step 3: Apply bi-directional z-score normalization

After hemispheric pooling, the correspondence matrix was represented with seven cortical systems as rows and cerebellar or subcortical networks as columns. This matrix was then normalized in two directions. First, column-wise z-scoring was applied within each cerebellar or subcortical network across the seven cortical systems, identifying the cortical system with the strongest relative correspondence for that network and reducing the influence of globally high or low cortical connectivity. Second, row-wise z-scoring was applied within each cortical system across cerebellar or subcortical networks, identifying noncortical networks with selectively strong correspondence to that system and reducing bias from cortical systems that were broadly connected to many noncortical networks. Together, the bi-directional z-score normalization made the assignment depend on relative selectivity rather than raw FC strength. This was particularly important in early life, when broad cortico-cerebellar and cortico-subcortical connectivity may reflect ongoing developmental segregation and integration rather than mature one-to-one functional correspondence.

#### Step 4: Assign cortical-system and cortical-family label

After normalization, each cerebellar or subcortical network was assigned to the cortical system with the highest adjusted correspondence score. This yielded the seven-system label for that network in the current bootstrap iteration. Fine-scale cortical-family assignment, i.e., 22-network, was then performed within the assigned cortical system. For each cerebellar or subcortical network, candidate cortical families were restricted to those belonging to its assigned seven-system class. The same pooling and normalization logic was then used to identify the cortical functional family with the strongest adjusted correspondence. This hierarchical strategy ensured that fine-scale family assignment remained consistent with the broader cortical-system organization.

#### Step 5: Summarize bootstrap statistics

The full assignment procedure was repeated across 1,000 bootstrap iterations. For each cerebellar or subcortical network, we computed the proportion of iterations in which it was assigned to each cortical system. The dominant cortical-system label was defined as the system with the highest assignment probability.

For division-specific scaffold map visualization, colors were assigned to encode reproducible functional correspondence across structures rather than to independently label each parcellation. This correspondence-informed color scheme differs from structure-specific color settings, in which cortical, cerebellar and subcortical parcels would each receive unrelated colors and cross-structure alignment would not be visually interpretable. We first assigned unique colors to the 22 cortical functional families, which served as the reference color scaffold. Cerebellar networks were then matched to cortical families within their assigned seven-system class, and each cortical family was allowed to share its color with at most one cerebellar network. When multiple cerebellar networks mapped to the same cortical family, the network with the strongest correspondence score was retained, yielding 18 cerebral cortex-matched cerebellar networks and 11 residual cerebellar networks. The same constrained matching procedure was applied to subcortical networks, yielding 18 cerebral cortex-matched subcortical networks and 13 residual subcortical networks. To capture additional cross-structure correspondence not resolved by cortical matching, residual subcortical networks were next matched directly to residual cerebellar networks within the same seven-system class; 7 residual subcortical networks were assigned the same color as their strongest cerebellar counterpart. The final division-specific scaffold map (Fig. 1b) contained 39 colors: 15 shared across cortex, cerebellum and subcortex; 3 shared by cortex and cerebellum only; 3 shared by cortex and subcortex only; 7 shared by cerebellum and subcortex only; 1 specific to a cortical family; 4 specific to cerebellar networks; and 6 specific to subcortical networks.

#### Quantification of correspondence robustness and specificity

We next quantified the robustness and specificity of these cross-structure correspondences using the bootstrap outputs of the seven-system assignment procedure. For each noncortical network *p*, correspondence to the cortical scaffold was represented by the empirical distribution of bootstrap-derived assignments across the seven cortical systems. Stability was defined as the probability of the modal cortical-system assignment across bootstrap iterations. If *π_pk_* denotes the proportion of bootstrap iterations in which network *p* was assigned to cortical system *k*, then

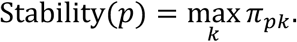

This metric quantifies how consistently a cerebellar or subcortical network mapped to the same cortical system across resampled datasets.

We quantified the distinctness of this correspondence using separability, defined as the median margin between the strongest and second-strongest cortical-system correspondence scores across bootstrap iterations. If 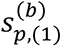 and 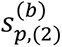 denote the highest and second-highest correspondence scores for network *p* in bootstrap iteration *b*, respectively, then

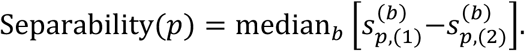

Whereas stability captures across-bootstrap consistency of the dominant cortical-system label, separability captures within-bootstrap distinctness of that label relative to its nearest alternative. To identify networks with weakly specified cortical correspondence, we combined stability and separability into a composite specificity score. Each metric was converted to its percentile rank across networks, and the final score was defined as

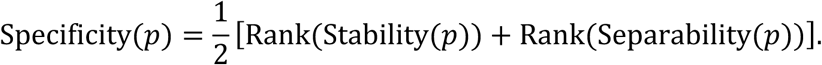

Networks with low composite scores were designated low-specificity networks, indicating both unstable cortical-system assignment and weak separation from competing alternatives.

To summarize ambiguity structure at the systems level, we further computed a system-admixture matrix from the top-ranked and second-ranked cortical-system assignments across all networks and bootstrap iterations. For each network in each bootstrap iteration, we recorded the dominant and runner-up cortical-system labels, accumulated their co-occurrence counts into a 7 × 7 matrix, and row-normalized the matrix to estimate the conditional probability of each second-choice system given the top-choice system,

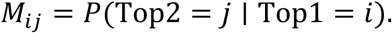

This matrix summarizes which cortical systems most frequently competed in explaining the correspondence profiles of cerebellar and subcortical networks, providing a systems-level description of overlap beyond network-specific measures.

### Whole-brain functional connectivity analyses

#### Functional connectivity matrix visualization

Whole-brain and structure-specific functional connectivity (FC) matrices were assembled from precomputed mean FC blocks spanning cerebral cortex, subcortex and cerebellum. For whole-brain visualization, blocks were concatenated in the order cerebral cortex, subcortex and cerebellum after parcel reordering to highlight mesoscale organization. Cortical parcels were grouped by the seven canonical functional systems within each hemisphere, subcortical parcels by major anatomical class, and cerebellar parcels by seven cortex-referenced functional groups. Matrices were displayed in a split-triangle format, with the lower triangle showing raw FC and the upper triangle showing a thresholded version. Thresholding was applied separately within each block for the whole-brain matrix and to the full matrix for within-structure panels; entries below the 60^th^ percentile of absolute FC magnitude were masked in gray, whereas the diagonal was retained from the raw matrix. FC values were clipped to fixed display ranges for visualization.

#### Force-directed whole-brain connectome layout

For graph visualization^59^, bilateral cortical and subcortical FC matrices were pooled across homologous left-right parcels using the same softmax pooling operator defined above (*β* = 1.5), yielding 22 cortical parcels and 31 subcortical parcels; the 29 cerebellar parcels were retained. This smooth max-like operator preserves strong unilateral contributions while avoiding attenuation from arithmetic averaging. Graph construction was based on absolute FC magnitude so that edge retention and node prominence reflected coupling strength rather than sign. Parcel-level matrices were then collapsed to 7 cortical systems, 7 cerebellar systems and 6 subcortical classes to define a macro-scale scaffold.

To reduce differences in dynamic range across FC blocks, each block was normalized by the 95^th^ percentile of its non-zero absolute values. System-level edges were then sparsified separately by connection class, retaining the strongest normalized fraction within each block (cortex-cortex, 35%; cortex-subcortex, 28%; cortex-cerebellum, 28%; cerebellum-subcortex, 25%) to account for differences in edge density across blocks while preserving comparable visual interpretability of within- and cross-structure organization, together with a minimum normalized cutoff of 0.06. Node positions were estimated using a weighted Fruchterman-Reingold spring layout implemented in NetworkX^60^ (spring_layout; *k* = 1.35, 1,400 iterations, seed = 1), with layout weights defined as ∣ FC ∣^0.B^. System node size was scaled by raw absolute connectivity strength aggregated across the corresponding FC blocks.

Parcel-level whole-brain connectomes were overlaid onto this scaffold by placing parcels as local satellites around the position of their parent cortical system, cerebellar network or subcortical class. After the same blockwise normalization, cross-structure parcel edges were selected by retaining, for each destination parcel, the two strongest cortex-to-subcortex edges, the two strongest cortex-to-cerebellum edges and the single strongest cerebellum-to-subcortex edge, subject to a minimum normalized cutoff of 0.12. Parcel node size was proportional to node strength, defined as the sum of retained raw absolute FC magnitudes across incident edges.

For the within-cortex, within-subcortex and within-cerebellum panels, parcels were displayed separately by structure while preserving the scaffold coordinates, identities and color assignments used in the whole-brain parcel panel. Within each structure, a new undirected graph was constructed from the corresponding square FC matrix by linking each parcel to its four strongest positive within-structure neighbors, with duplicate edges removed. Node size was rescaled according to within-structure node strength, defined as the sum of retained incident edge weights. Thus, parcel positions were held constant across panels, whereas edges and node strengths were re-estimated separately for cerebral cortex, subcortex and cerebellum.

### Developmental trajectory analysis of whole-brain functional connectivity

#### Modeling developmental trajectories of functional connectivity

Developmental changes in each FC feature were estimated using a generalized additive mixed model (GAMM)^61^. For each feature, FC was modeled as a smooth function of scan age with a participant-level random effect to account for repeated longitudinal measurements: *y* ∼ *s*(Scan Age, *bs* = *cr*, *k* = 8) + *s*(ID, *bs* = “*re*”). The age effect was estimated using a cubic regression spline basis, and model parameters were estimated by restricted maximum likelihood. Features with fewer than 30 valid observations or fewer than three unique participants were not fitted. Population-level developmental trajectories were obtained by predicting across the observed age range on a dense grid of 801 points while excluding the participant-level random effect. Pointwise 95% confidence intervals were calculated from prediction standard errors as the fitted value ±1.96 standard errors. In trajectory plots, gray dots indicate individual observations, solid curves denote fitted population-level trajectories and shaded bands indicate pointwise 95% confidence intervals. Sex was not included as a covariate in these primary models. As supplementary analyses, we fitted sex-stratified GAMMs including a main effect of sex, sex-specific smooths of age and a subject-level random effect (*y* ∼ Sex + *s*(ScanAgeY, by = Sex) + *s*(ID, *bs* = “*re*”)), and present sex-specific developmental trajectories of within-network FC, the top 40 connections showing sex differences and the top 40 connections showing the largest age-related differences in Supplementary Figs. 3-5.

#### System-level visualization of developmental FC change patterns

To summarize developmental FC organization across 0-6 years, fitted edge-wise GAMM trajectories were restricted to the 0-6-year interval, aligned to a common age axis and grouped according to the anatomical or system identities of their endpoints. Trajectories within each matrix cell were averaged pointwise to obtain a mean developmental profile. Each profile was then classified as consistently positive, consistently negative, positive-to-negative or negative-to-positive according to its sign pattern across age, and its mean FC over 0-6 years was computed. Summary matrices were displayed in lower-triangular form, with tile color indicating trajectory class and circle size indicating the absolute magnitude of mean FC. We used both a unified 21-node representation, in which cortex, subcortex and cerebellum were each summarized into seven matched large-scale systems, and a complementary 20-node representation, in which the subcortex was resolved into six anatomical classes while cortex and cerebellum retained the same seven-system organization.

#### Distribution of developmental trajectory patterns across connection types

To examine whether characteristic developmental FC trajectories were preferentially represented in specific anatomical connection classes, fitted edge-wise GAMM trajectories were summarized using a rule-guided pattern classification. We first explored unsupervised trajectory clustering, but the resulting solutions were sensitive to clustering settings and often separated curves by amplitude rather than by interpretable temporal shape. We therefore inspected the fitted trajectories, identified the dominant recurring profiles, and defined eight developmental trajectory patterns: early decline with late rebound, low-amplitude transient increase followed by decline, early high-positive decline, transient mid-childhood peak followed by decline, gradual monotonic increase, high-positive U-shaped trajectory with late rebound, stable positive trajectory and gradual monotonic decline. This classification was used as a descriptive framework to summarize large-scale developmental organization rather than as an independent inferential procedure. Connection type was defined directly from feature labels and grouped into six classes: cortex-cortex (CC), cortex-cerebellum (CI), cortex-subcortex (CS), cerebellum-cerebellum (II), subcortex-subcortex (SS) and cerebellum-subcortex (IS). For each trajectory pattern, we counted the number of contributing edges from each connection class and calculated its proportion among all analyzed edges. Conversely, for each connection class, we quantified the within-class percentage distribution across the eight trajectory patterns. These summaries were visualized using complementary stacked bar plots showing trajectory-pattern counts stratified by connection type and within-class percentage distributions across patterns.

#### Turning-point timing analysis

To quantify the timing of major developmental inflections, we analyzed the fitted edge-wise GAMM trajectories and identified turning points from sign changes in the first derivative^62^. Positive-to-negative transitions were classified as peaks and negative-to-positive transitions as valleys. A small derivative tolerance was used to suppress numerical fluctuations near zero, and plateau-like extrema were classified according to the direction of the surrounding slope. For each trajectory, only the first detected turning point was retained to capture the earliest major developmental reorganization of each FC feature and to limit the influence of later secondary undulations or smoothing-related fluctuations. To summarize timing across the connectome, we estimated smoothed density curves for first turning-point ages over 0-72 months. Separate densities were computed for peak and valley events and weighted by their relative frequencies among all trajectories with an identified turning point. An overall density was also estimated from all first turning-point ages combined. These distributions summarize both the timing and prevalence of early developmental inflections across FC trajectories.

### Prediction of age, sex, and cognition from whole-brain functional connectivity

Prediction analyses were based on whole-brain functional connectivity (FC) features derived from a common 7-network representation spanning cerebral cortex, subcortex and cerebellum. Parcel-level connectivity within each structure was collapsed into seven matched large-scale functional systems, yielding a 21 × 21 whole-brain FC matrix. From this matrix, we defined six FC classes capturing the principal within- and between-structure components of the whole-brain scaffold: cortex-cortex (CC), cerebellum-cerebellum (II), subcortex-subcortex (SS), cortex-cerebellum (CI), cortex-subcortex (CS) and cerebellum-subcortex (IS). Within-structure classes were vectorized using the upper triangle including the diagonal, whereas between-structure classes were vectorized using all pairwise entries, resulting in 231 FC features per sample. Mean framewise displacement was included as a covariate to reduce residual motion-related effects.

We examined three complementary prediction targets. Age prediction tested whether the whole-brain FC scaffold captured orderly developmental variation across early life. Sex prediction tested whether the whole-brain functional scaffold contained reproducible connectivity patterns associated with sex. Prospective cognitive prediction used FC measured at the imaging visit to predict cognitive performance assessed 6-12 months later, thereby testing whether early whole-brain functional organization was informative of subsequent developmental outcome. Final cognitive analyses focused on the Early Learning Composite and five subdomain scores: expressive language, fine motor, gross motor, receptive language and visual reception. The interval between imaging and cognitive assessment was included as an additional covariate for sex and cognitive prediction, but not for age prediction.

Model evaluation used repeated subject-level cross-validation (5 repeats of 5 folds) to ensure that no participant contributed data to both training and test sets within a fold. Continuous outcomes were evaluated using grouped subject-level folds, whereas sex classification used subject-level stratified folds. To assess whether predictive information was preferentially concentrated in specific portions of the whole-brain scaffold, we evaluated all 63 non-empty combinations of the six FC classes. Within each training fold, features were standardized and reduced using univariate feature selection (SelectKBest^63^), retaining 50% of available FC features in the selected FC-class combination (capped at 1,024 features), with all features retained when a combination contained 50 or fewer features. Feature ranking used regression (f_regression) for continuous outcomes and classification (f_classif) for sex classification. Models were fixed by target type: Lasso regression^64^ for age, random forest regression^65^ for cognitive outcomes and radial-basis-function support vector classification^66^ for sex. Predictive performance was quantified primarily by Pearson correlation for continuous outcomes and area under the receiver operating characteristic curve^67^ for sex classification.

To identify the components of the whole-brain FC scaffold contributing most strongly to prediction, feature importance was estimated within the same repeated subject-level cross-validation framework. For each target and each of the 63 non-empty FC-class combinations, importance was computed after univariate feature selection and then mapped back to the full 231-feature FC space, with unselected features assigned zero importance. For age prediction, importance was defined as the absolute value of retained Lasso coefficients; for cognitive prediction, as random forest feature importance; and for sex classification, as held-out permutation importance. Importance values were accumulated across folds, repeats and FC-class combinations and then averaged to obtain a target-specific importance score for each FC feature. These connection-level importance estimates were subsequently collapsed into the six FC classes to quantify their broader contributions to prediction.

Results were summarized at three complementary levels. First, predictive performance was evaluated both for the optimal FC-class combination for each target and across all FC-class combinations. Second, the most contributive FC features were visualized as circular connectograms after mapping each feature to its structural assignment and 7-network identity. Third, FC-class contributions were summarized by quantifying how often each FC class appeared among the top-performing combinations and by normalizing the summed importance of each FC class within each target to compare their relative contributions across prediction domains.

## Acknowledgments

We thank the participating children and families and the UNC/UMN Baby Connectome Project Consortium for their contributions to this study. This work was supported in part by the National Institutes of Health grants HD119560, AG075582, NS128534, EB037388 and NS135574.

## Author contributions

D.H. conceived the study, developed the methodology, led the analyses, interpreted the results, prepared the figures and wrote the manuscript. J.C. contributed substantially to method development, implementation, data analysis and manuscript preparation. K.H. contributed to algorithm design, software development, preprocessing and diffusion MRI analyses. W.Y. and Y.S. performed data preprocessing, quality control and data preparation. J.L. contributed to visualization and figure preparation. S.-C.H. and J.R.C. contributed neuroscience expertise and interpretation. L.W. supervised data preprocessing and contributed to methodological discussions and interpretation. W.L. and G.L. jointly supervised the study, contributed to study design, methodological development and scientific interpretation and critically revised the manuscript. All authors reviewed and approved the manuscript.

## Data availability

The UNC/UMN Baby Connectome Project (BCP) and Developing Human Connectome Project (dHCP) data are available through the NIMH Data Archive (NDA), subject to NDA registration and data-access procedures. The Pixar dataset is publicly available through OpenNeuro (http://openneuro.org/datasets/ds000228/). The Calgary Preschool MRI Dataset is publicly available through the Open Science Framework (https://osf.io/axz5r/). The MAP dataset is available from the corresponding authors upon reasonable request.

## Code availability

This study used openly available software and code. The Early Life Whole-Brain parcellation, data processing, and analysis codes are available at: https://www.nitrc.org/projects/unc_elf.

