## Supplemental Data 1 for "Division-Specific Organization of a Shared Functional Scaffold in the Early-Life Human Brain"

Dan Hu *et al.*

**This PDF file includes:**

Extended Data Fig. 1 to Fig. 9

Supplementary Fig. 1 to Fig. 5

Supplementary Table S1 to Table S2

### Extended Data

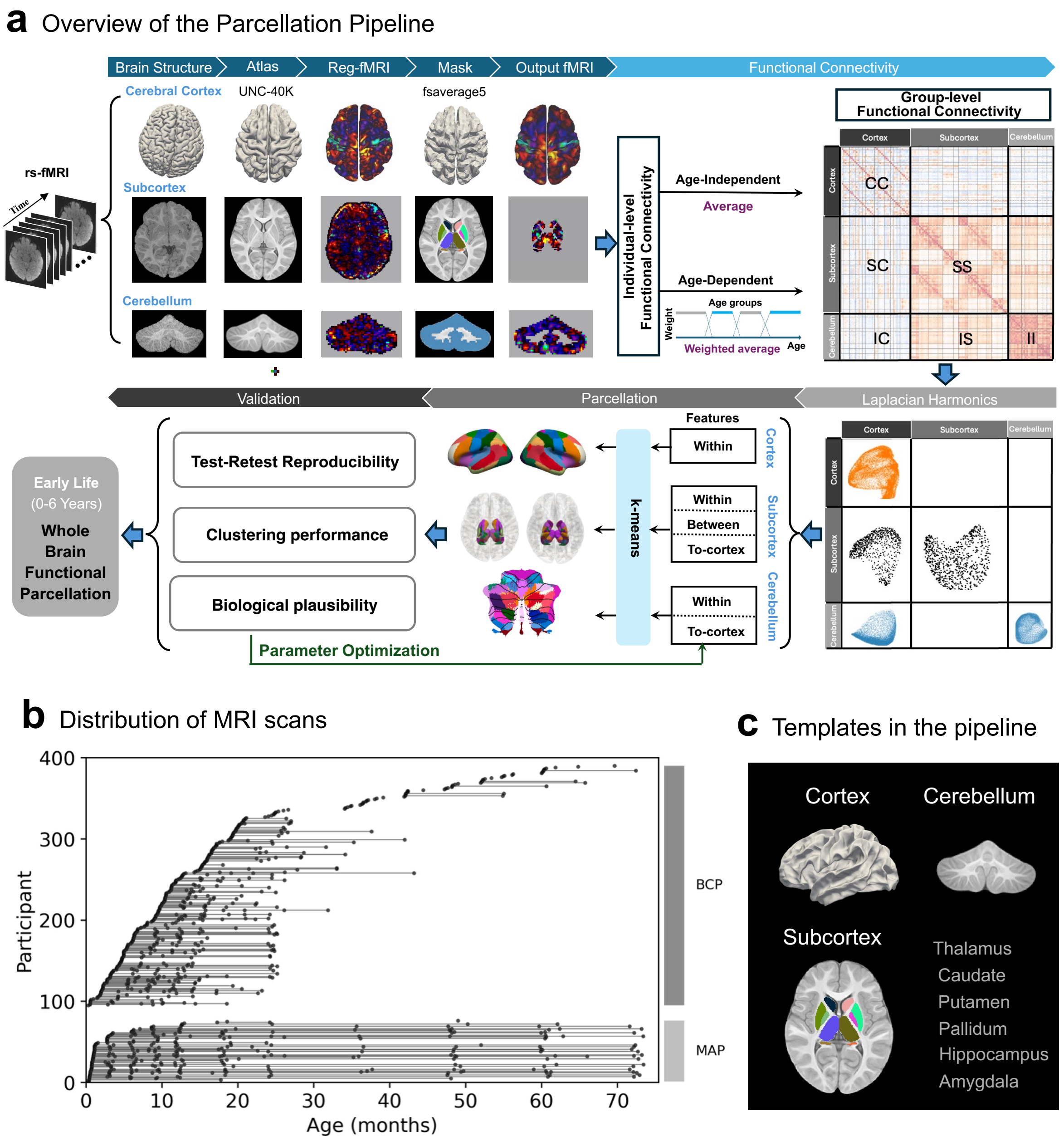

**Extended Data Figure 1**. Workflow of the unified early-life whole-brain functional parcellation framework. **a**, Overview of the parcellation pipeline. Resting-state fMRI data acquired from children aged 0-6 years were processed separately for the cerebral cortex, subcortex and cerebellum using structure-appropriate atlases, registrations and masks. Structure-specific time series were extracted and used to compute individual-level functional connectivity matrices. These matrices were aggregated either across all scans to generate age-independent group-level connectivity or within overlapping age windows to generate age-dependent connectivity. Group-level connectivity matrices included both within-structure and cross-structure interactions and were transformed into Laplacian harmonic representations. Structure-specific feature sets were then constructed for parcellation: intra-cortical features for the cerebral cortex, within- and between-subcortical plus subcortico-cortical features for the subcortex, and within-cerebellar plus cerebello-cortical features for the cerebellum. K-means clustering was applied within each division to generate candidate parcellations. Parameter selection was guided by test-retest reproducibility, clustering performance and biological interpretability, yielding the Early-Life Whole-Brain Functional Parcellation (UNC-ELF). **b**, Distribution of MRI scans across participants and age in the BCP and MAP datasets. Each horizontal line denotes the longitudinal scanning profile of one participant, illustrating the age coverage and sampling density across early life. **c**, Structure-specific templates used in the pipeline, including cortical, subcortical and cerebellar templates for feature extraction and parcellation.

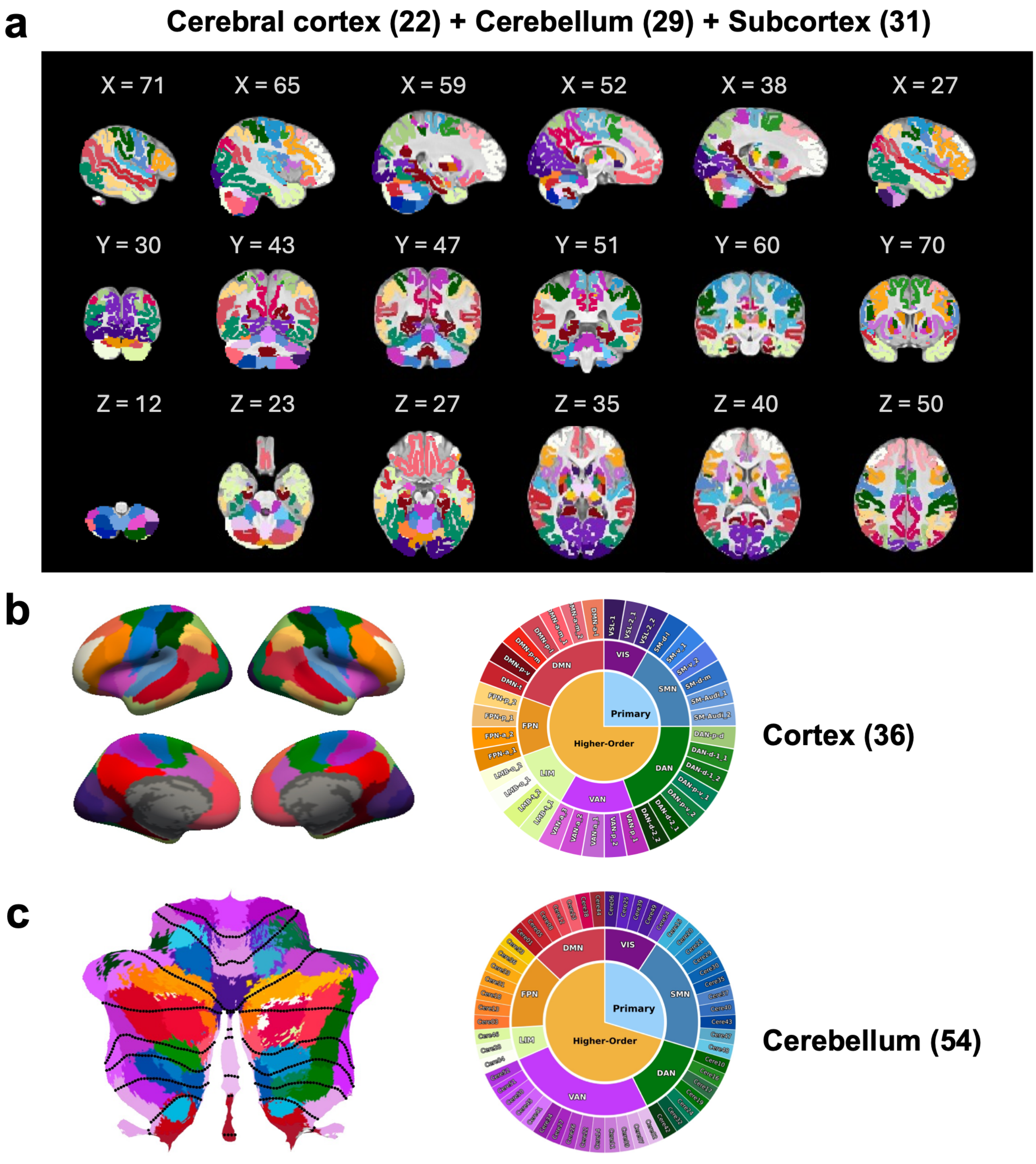

**Extended Data Figure 2**. Volumetric rendering of the early-life whole-brain parcellation (UNC-ELF) and alternative cortical and cerebellar resolutions. **a**, Main-resolution whole-brain parcellation displayed in the 24-month UNC volumetric template. The surface-derived 22-network cerebral cortical parcellation was projected into volumetric space and visualized together with the 29-network cerebellar and 31-parcel subcortical parcellations across sagittal, coronal and axial slices. This rendering illustrates the spatial coverage of the principal parcellation used in the main analyses across cerebral cortex, cerebellum and subcortex. **b**, Higher-resolution cerebral cortical parcellation comprising 36 networks, shown on cortical surface views and summarized with a nested network wheel. This solution provides a finer-scale cortical representation complementary to the principal 22-network scheme. **c**, Higher-resolution cerebellar parcellation comprising 54 networks, shown on the cerebellar flatmap and summarized with a nested network wheel. This solution provides a finer-scale cerebellar representation complementary to the principal 29-network scheme. For visualization of the higher-resolution cortical and cerebellar solutions, colors were assigned according to correspondence with the seven-network organization. Cortical correspondence was determined by Dice overlap with the seven-network cortical solution derived from the 22-network parcellation, whereas cerebellar correspondence was assigned using cortex-referenced functional connectivity as described in the Online Methods.

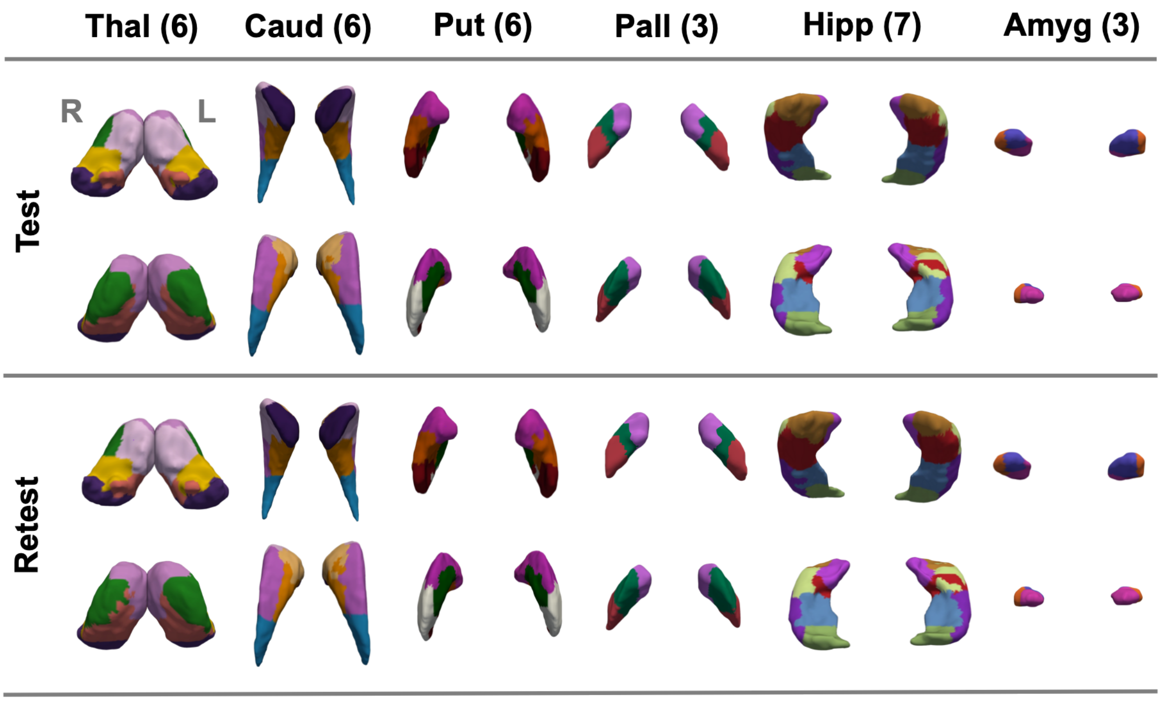

**Extended Data Figure 3.** Surface renderings of subcortical parcellations derived independently from test and retest data are shown for the selected resolutions identified in Fig. 4c. Bilateral thalamus, caudate, putamen, pallidum, hippocampus and amygdala show highly consistent parcel topology across independent splits, supporting the reproducibility of the subcortical parcellation. Two views are shown for each structure to illustrate spatial correspondence across the full extent of each nucleus. For the hippocampus, minor local boundary differences are observed when comparing the split-specific test–retest solutions with the final solution estimated from all available scans. This is expected because the test, retest and final parcellations were estimated from different data subsets, and local boundaries in small, curved structures such as the hippocampus can shift slightly when sample composition and signal averaging change. These small local differences do not alter the overall parcel topology or the reproducibility pattern quantified by the test–retest Dice analyses in Fig. 4c.

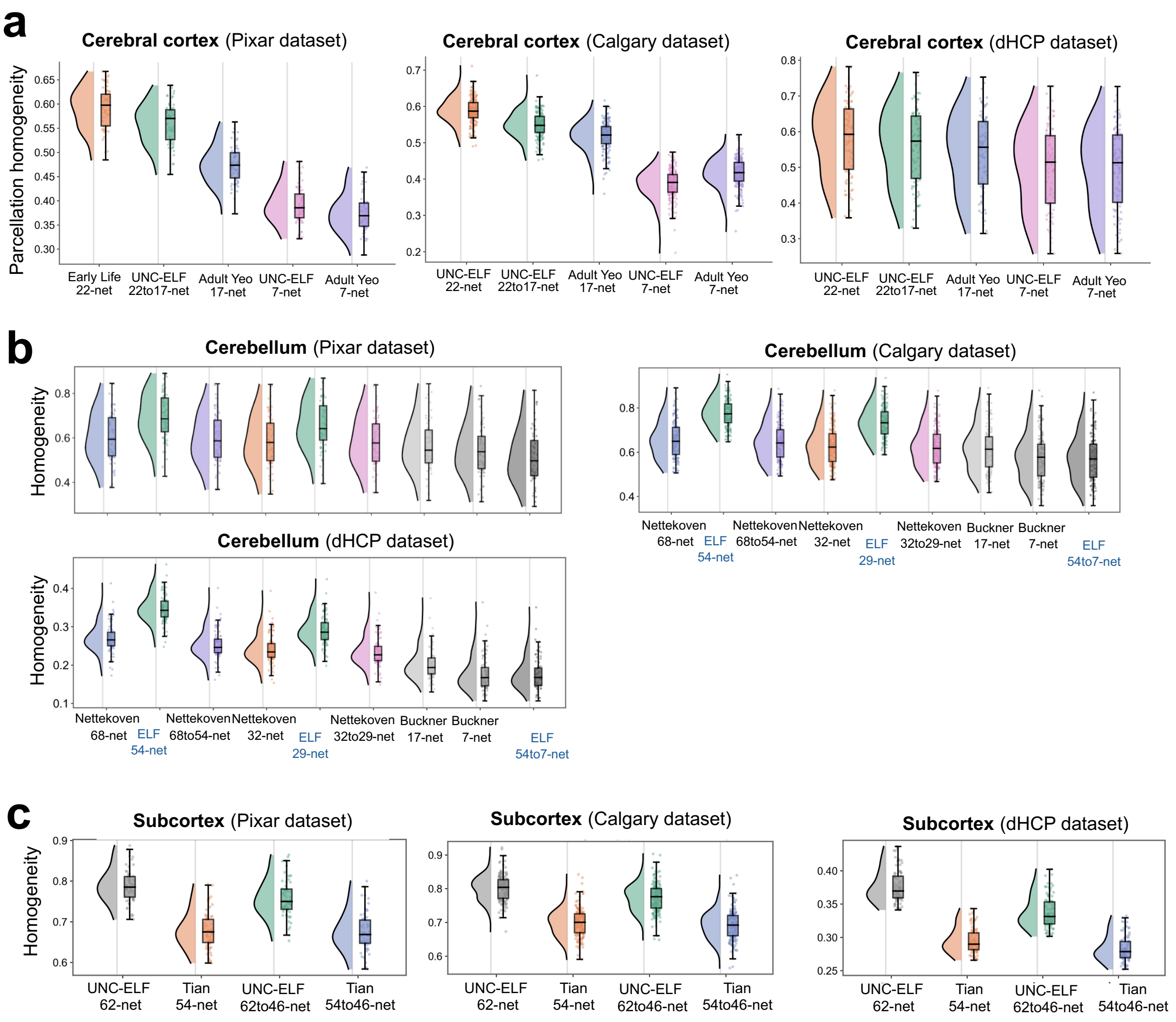

**Extended Data Figure 4**. Functional homogeneity of the Early-Life Whole-Brain Functional Parcellation (UNC-ELF) in independent external datasets. Functional homogeneity was evaluated in three external early life cohorts (Pixar, Calgary and dHCP) to assess the generalizability of the UNC-ELF beyond the discovery data. **a**, Cortical homogeneity comparisons across external datasets. The UNC-ELF 22-network and 22-to-17-network cortical parcellations showed higher within-parcel functional homogeneity than adult Yeo reference parcellations across external cohorts. At the coarser seven-network scale, homogeneity was more comparable to Yeo-7, consistent with the reduced sensitivity of broad canonical systems to developmental boundary differences and with the use of the seven-network representation as a shared cross-structure reference. Together, these results support the developmental relevance and external robustness of the cortical solution, particularly at finer functional resolutions. **b**, Cerebellar homogeneity comparisons across external datasets. UNC-ELF showed higher within-parcel functional homogeneity than adult-derived cerebellar reference maps across multiple matched-resolution comparisons, including Nettekoven- and Buckner-based schemes, supporting improved fit to early-life cerebellar functional organization. **c**, Subcortical homogeneity comparisons across external datasets. UNC-ELF subcortical parcellation showed higher functional homogeneity than the adult Tian reference parcellation at both the native whole-subcortex resolution and after harmonization to a matched 46-label representation. For simplicity, only whole-subcortex results are shown here rather than nucleus-specific comparisons. Together, these results indicate that the improved functional precision of UNC-ELF generalizes across independent early-life samples. For all box plots: center line, median; box limits, upper and lower quartiles; whiskers, 1.5× interquartile range; points, individual subjects.

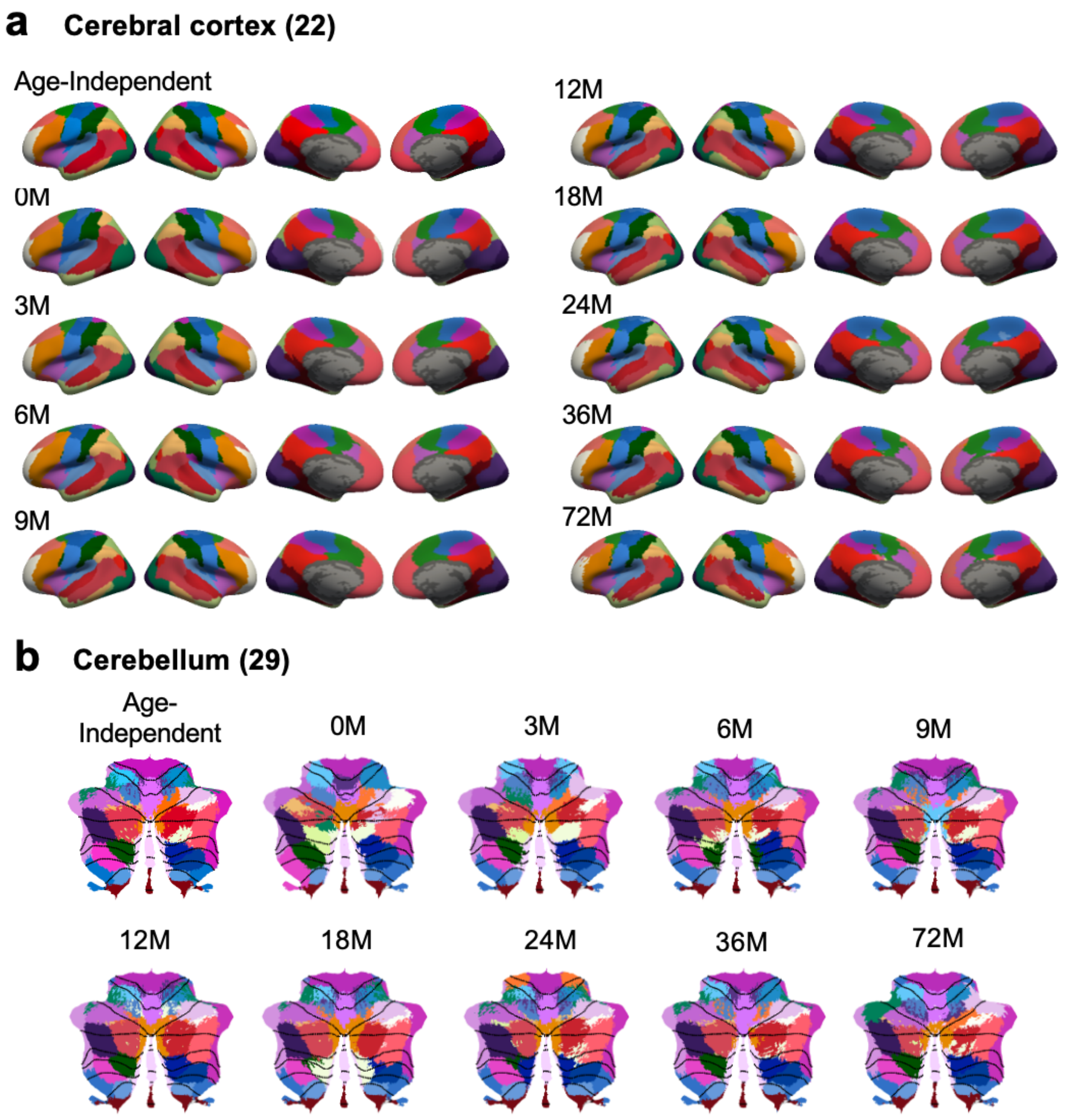

**Extended Data Figure 5**. Age-dependent cortical and cerebellar parcellations at the principal resolution. Age-dependent parcellations were estimated separately across developmental age groups and compared with the age-independent solution at the principal resolutions used in the main analyses. **a**, Cerebral cortical parcellations at the 22-network resolution, shown on inflated cortical surface views for the age-independent solution and each developmental age group. **b**, Cerebellar parcellations at the 29-network resolution, shown on the cerebellar flatmap for the age-independent solution and each developmental age group. Across both the cerebral cortex and cerebellum, the large-scale parcellation pattern is broadly preserved across age groups, with developmental differences expressed mainly as local boundary variation rather than wholesale reconfiguration of the functional organization. These age-dependent results support the presence of a broadly preserved macroscale scaffold across early life, while indicating that developmental effects are reflected primarily in finer spatial variation within this shared organizational framework. M: months.

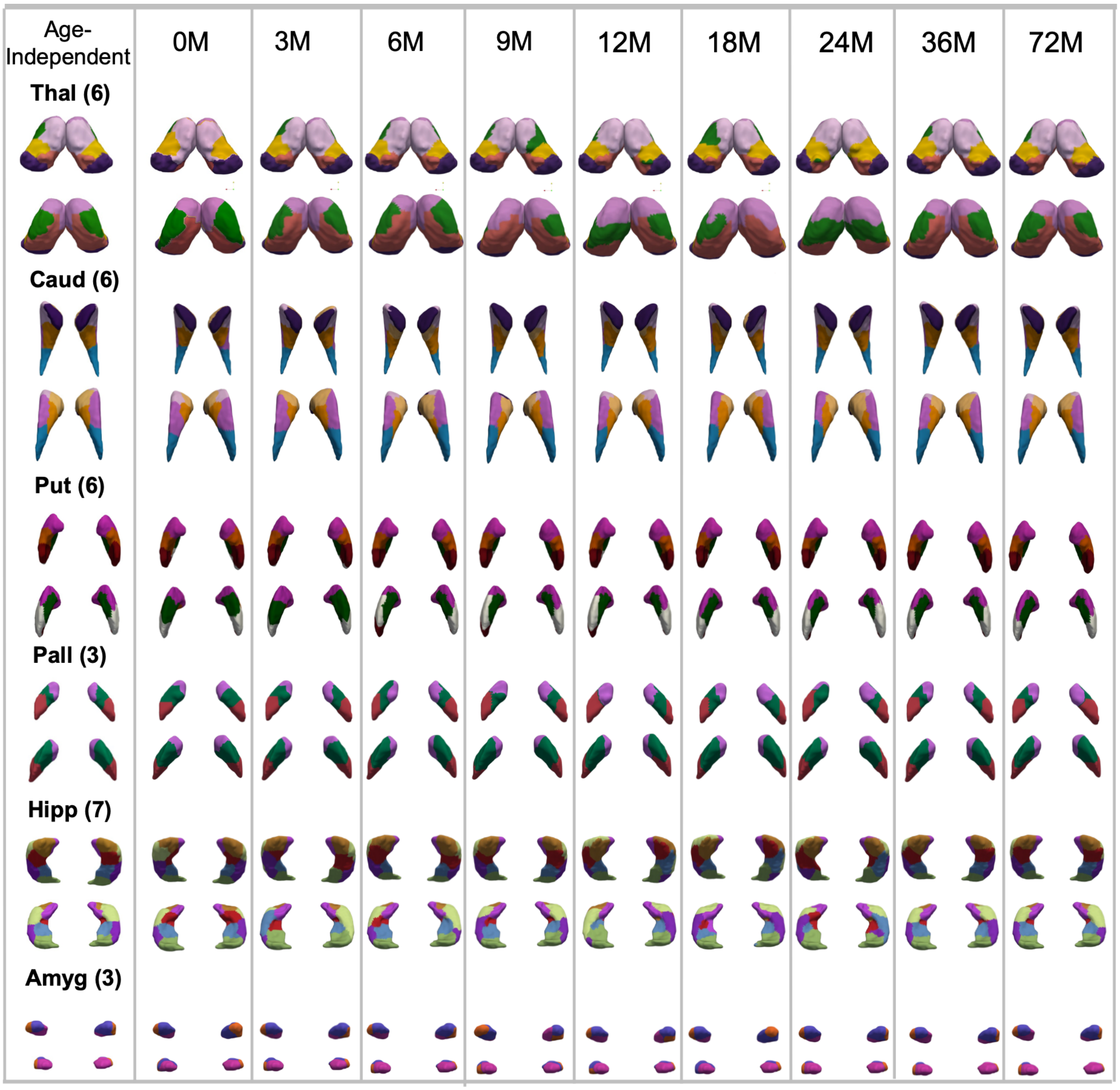

**Extended Data Figure 6**. Age-dependent subcortical parcellations at the principal resolution. Age-dependent subcortical parcellations were estimated separately across developmental age groups and compared with the age-independent solution at the same resolutions used in the main analyses. Bilateral thalamus (6 parcels), caudate (6 parcels), putamen (6 parcels), pallidum (3 parcels), hippocampus (7 parcels) and amygdala (3 parcels) are shown across age groups in two views for each structure. Across nuclei, the overall subcortical parcellation pattern is broadly preserved across development, with age-related differences expressed mainly as local boundary variation rather than large-scale reconfiguration. These results are consistent with the age-dependent findings in the cerebral cortex and cerebellum, and support the presence of a stable macroscale subcortical functional scaffold across early life.

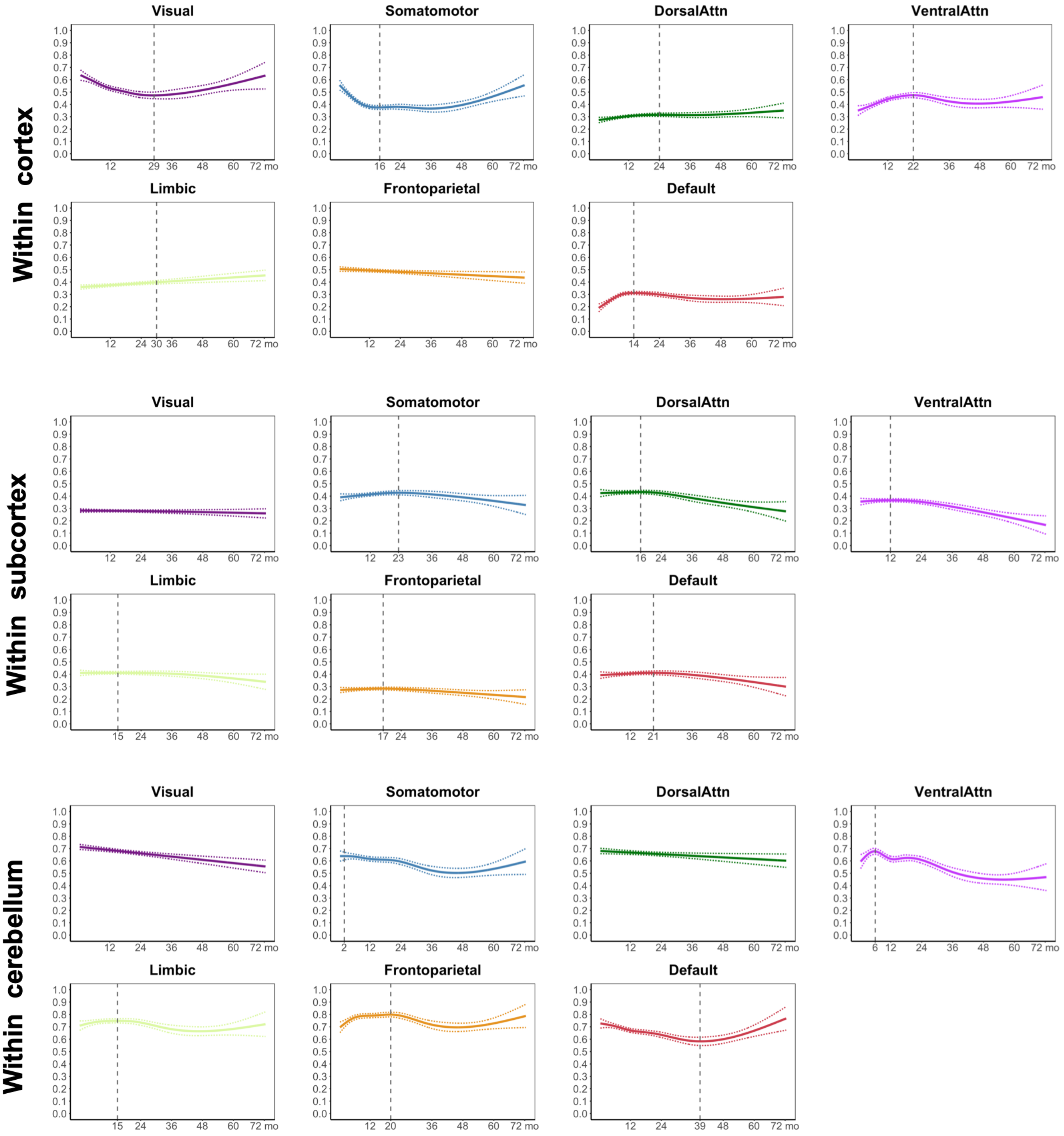

**Extended Data Figure 7.** Structure-specific developmental trajectories of within-network functional connectivity across early life. Developmental trajectories of within-network functional connectivity from birth to 6 years are shown for matched large-scale functional systems within the cortex, subcortex and cerebellum. Each panel shows the population-level trajectory fitted using a generalized additive mixed model (GAMM), with thin lines indicating 95% confidence intervals. Vertical dashed lines mark the first detected turning point of the fitted trajectory when present. Within-cortex trajectories show modest but heterogeneous developmental profiles across systems.

Within-subcortex trajectories are comparatively flatter, with weaker age-related variation. Within-cerebellum trajectories show larger amplitudes and greater nonlinearity, including declining, rebound-like and U-shaped profiles across sensory, motor and association systems. These patterns indicate that within-network maturation is structure-specific rather than uniform across the early-life whole-brain scaffold.

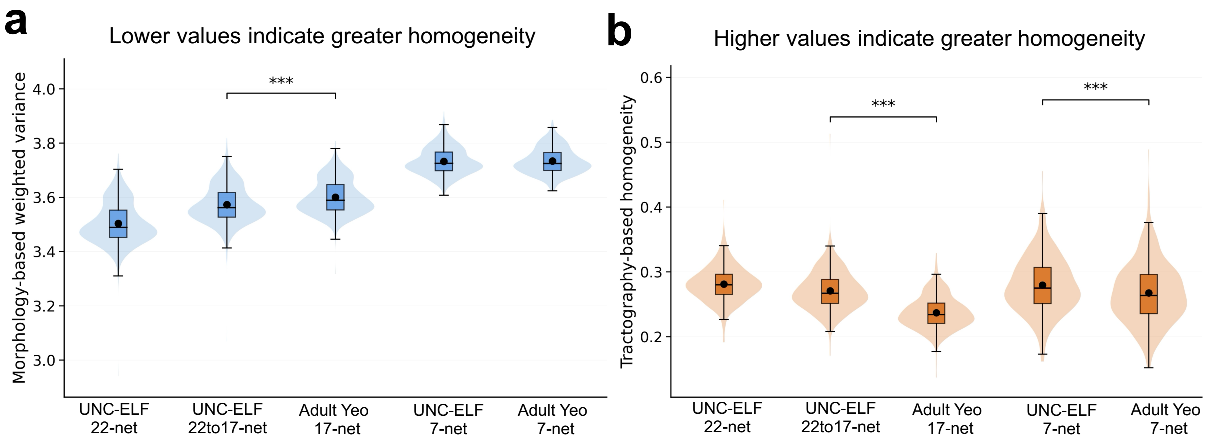

**Extended Data Figure 8**. Morphology- and tractography-based homogeneity of cortical parcellation schemes. **a**, Morphology-based homogeneity across cortical parcellation schemes. Violin plots show the distribution of subject-level morphology-based weighted variance, with embedded box plots indicating the interquartile range and squares indicating the mean. Morphological homogeneity was quantified by computing, within each parcel, the variance of vertex-wise morphological features, including sulcal depth, cortical thickness, surface area and curvature, after z-scoring across vertices within each subject. Parcel-level variances were then averaged across parcels weighted by parcel size. Lower values indicate greater within-parcel morphological homogeneity. **b**, Tractography-based homogeneity across the same cortical parcellation schemes. Tractography-based homogeneity was quantified as the within-parcel similarity of tractography-derived connectivity fingerprints, averaged across parcels weighted by parcel size. Higher values indicate greater within-parcel tractography-based homogeneity. Across both metrics, the UNC-ELF cortical parcellation at the principal resolution (22-net) and its derived lower-resolution representations (22→17-net and 22→7-net) were compared with adult Yeo-17 and Yeo-7 reference parcellations. At the 17-network scale, the UNC-ELF 22→17-net representation showed lower morphology-based weighted variance and higher tractography-based homogeneity than the adult Yeo-17 atlas, supporting improved structural coherence at finer cortical resolution. At the 7-network scale, performance was more comparable, consistent with the broad nature of canonical large-scale systems. (***, $p < 0.001$, paired *t*-test).

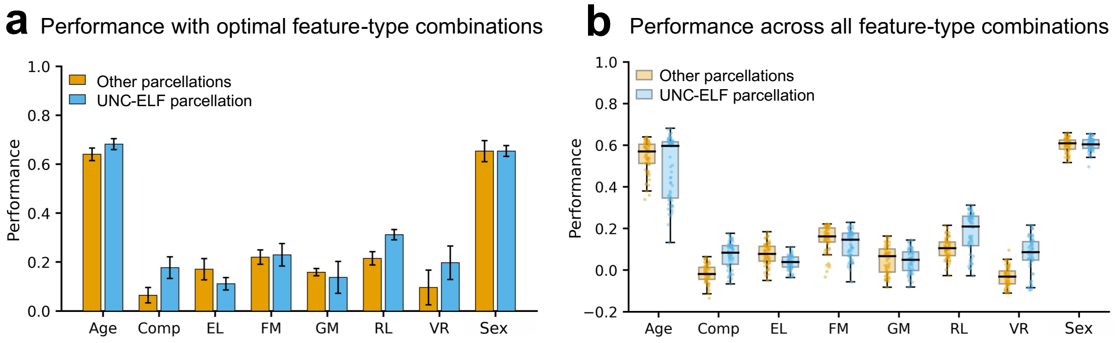

**Extended Data Figure 9**. Developmental and cognitive prediction comparison with an adult-derived whole-brain reference scheme. Prediction performance was compared between the Early-Life Whole-Brain Functional Parcellation (UNC-ELF) and an adult-derived whole-brain reference scheme composed of the Yeo 7-network cortical parcellation, the Buckner 7-network cerebellar parcellation and a 7-network subcortical parcellation obtained by merging the Tian-54 atlas according to overlap with the UNC-ELF 7-network solution. **a**, Optimal prediction performance for age at scan, sex and prospective cognitive outcomes assessed 6-12 months after scanning. Bar heights indicate the mean performance of the single best-performing combination of functional connectivity (FC) classes for each target. Performance is reported as Pearson correlation coefficient (r) for age and continuous cognitive measures, and area under the receiver operating characteristic curve (AUC) for sex classification. Error bars indicate s.d. across repeated cross-validation folds. **b**, Prediction performance across all 63 possible non-empty combinations of the six FC classes (cortex-cortex, cerebellum-cerebellum, subcortex-subcortex, cortex-cerebellum, cortex-subcortex, and cerebellum-subcortex). Box plots summarize the distribution of model performance across FC-class combinations for each target. Across targets, the UNC-ELF parcellation generally matched or exceeded the adult-derived reference scheme, supporting its utility for prediction of developmental state and prospective cognitive outcomes.

#### Supplementary Information

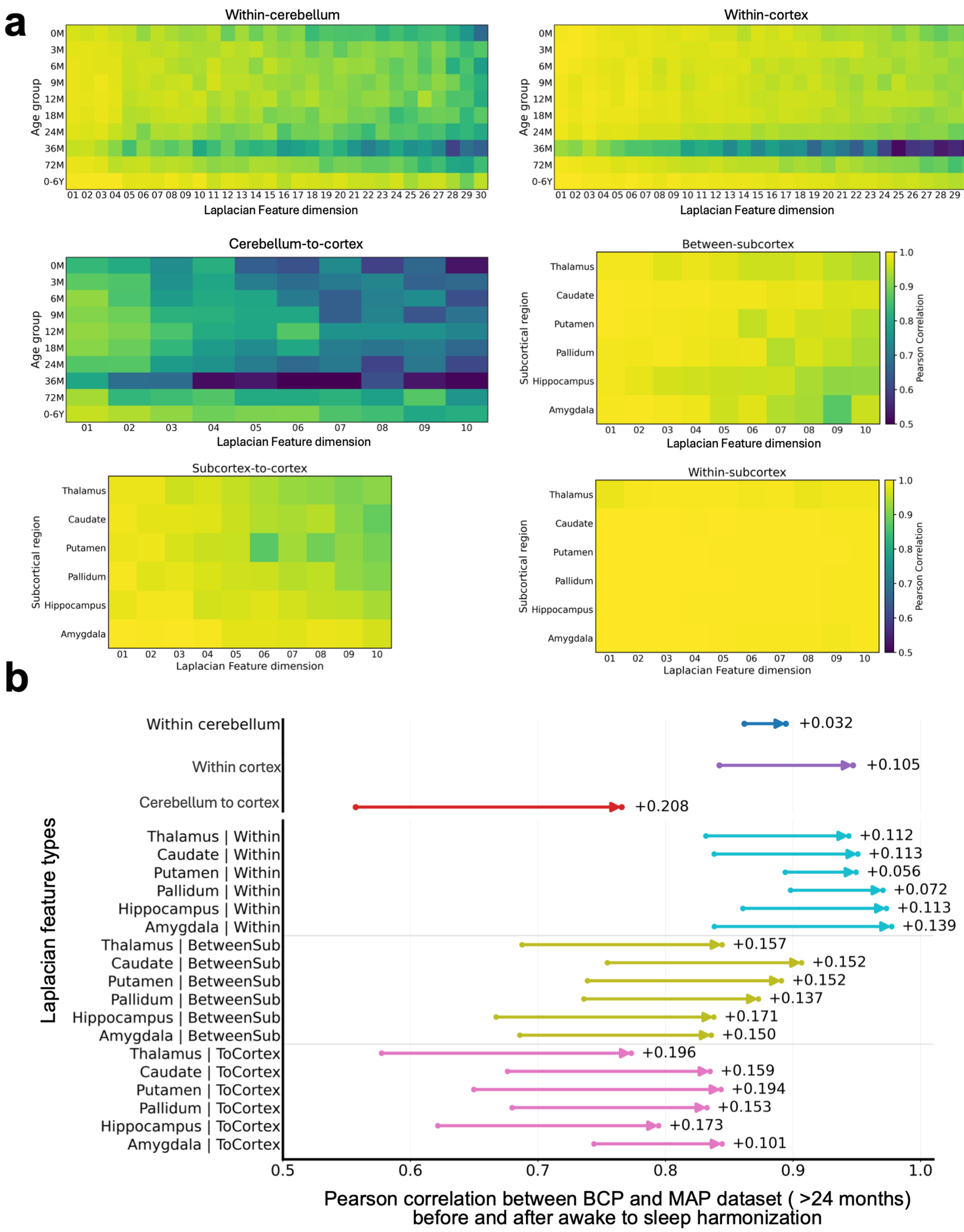

**Supplementary Figure 1**. Awake-to-sleep harmonization increases cross-dataset correspondence of Laplacian embedding features between BCP and MAP. **a**, Cross-dataset correspondence of

Laplacian embedding features derived from BCP and MAP before awake-to-sleep harmonization. Pearson correlations were computed between corresponding embedding dimensions across datasets. For whole-brain connectivity features, heat maps show BCP-MAP correspondence across age groups for within-cerebellum, within-cortex and cerebello-cortical embeddings. For subcortical features, heat maps show correspondence across subcortical regions for between-subcortex, subcortico-cortical and within-subcortex embeddings. Columns indicate leading Laplacian dimensions, and warmer colors indicate stronger cross-dataset agreement. Correspondence was generally higher for early embedding dimensions and within-structure features, whereas later dimensions and long-range cross-structure features showed lower agreement, suggesting that scan-state differences contributed to cross-dataset variability. **b**, Effect of awake-to-sleep harmonization on BCP-MAP agreement. Arrows connect mean Pearson correlations before and after harmonization for each feature set, with values at right indicating the absolute increase. The top three rows summarize whole-brain connectivity features computed across the first 30 embedding dimensions for age groups older than 24 months. The lower rows summarize subcortical features computed across the first 10 embedding dimensions and averaged across hemispheres and age groups older than 24 months. Awake-to-sleep harmonization increased cross-dataset agreement across all feature types, with the largest improvements observed for cerebello-cortical and subcortico-cortical embeddings.

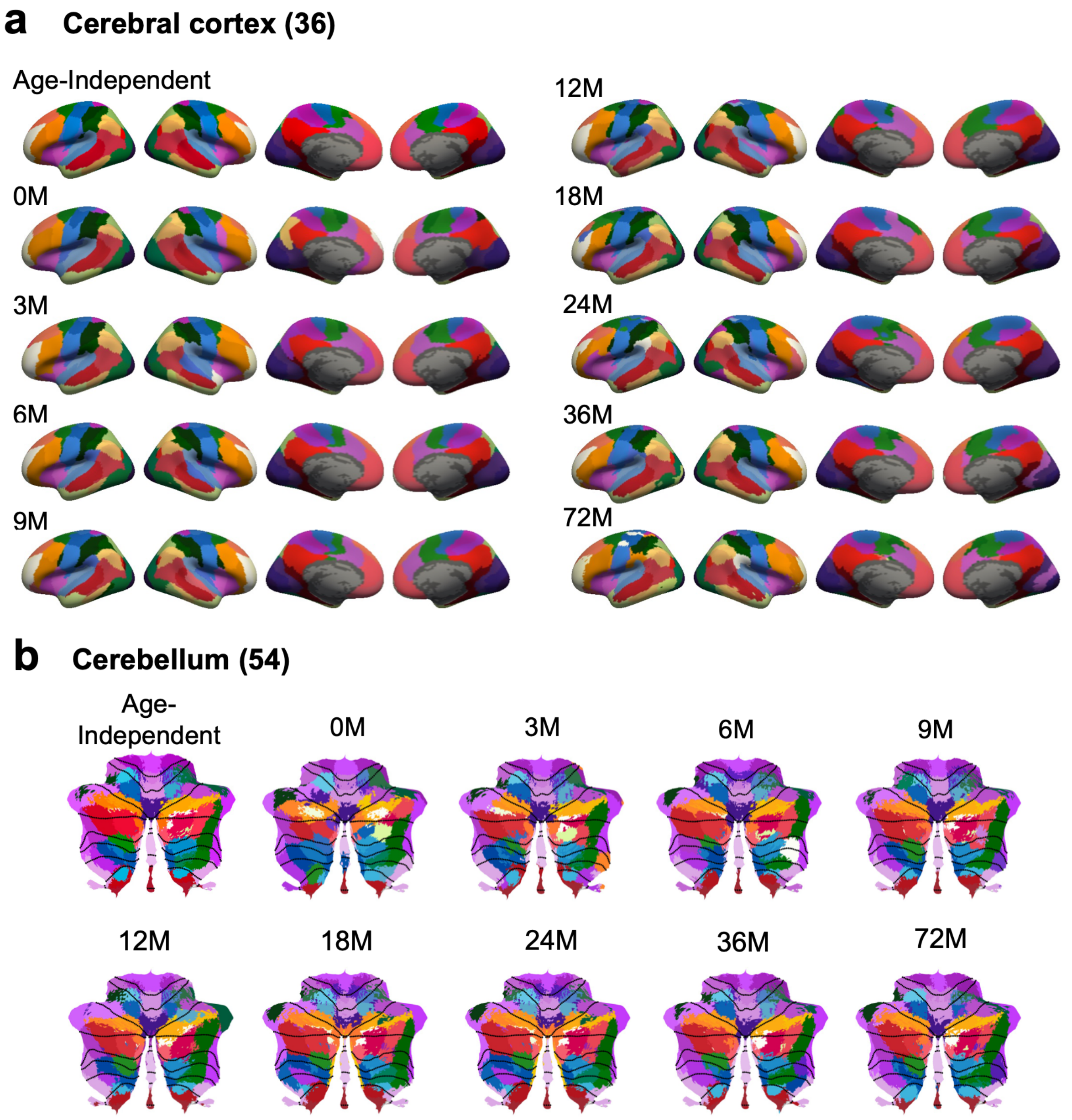

**Supplementary Figure 2**. Age-dependent cortical and cerebellar parcellations at higher resolution. Age-dependent parcellations were estimated separately across developmental age groups and compared with the age-independent solution at higher resolutions than those used in the main analyses, including the 36-network cerebral cortex and 54-network cerebellum. **a**, Cortical parcellations at the 36-network resolution, shown on inflated surface views for the age-independent solution and each developmental age group. **b**, Cerebellar parcellations at the 54-network resolution, shown on the cerebellar flatmap for the age-independent solution and each developmental age group. Across both structures, the overall large-scale organization is broadly preserved across development, with age-related differences expressed mainly as local boundary variation rather than major reconfiguration of the functional scaffold. These higher-resolution results further support the stability of the macroscale organizational framework across early life.

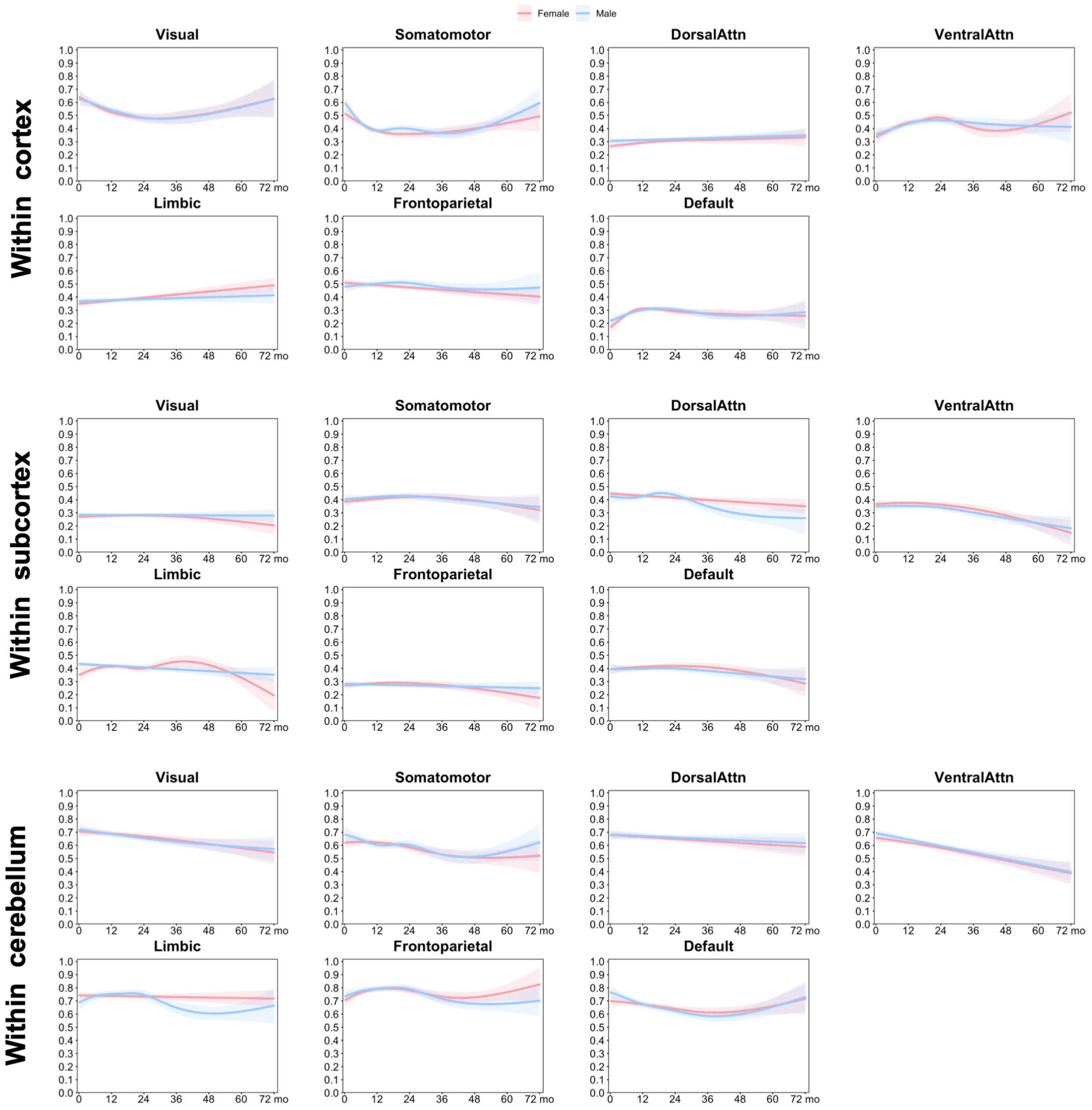

**Supplementary Figure 3**. Sex-specific developmental trajectories of within-network functional connectivity across early life. Sex-specific developmental trajectories of within-network functional connectivity are shown for females and males across the same structure-specific connectivity classes and large-scale functional systems presented in Extended Data Fig. 7. Trajectories were fitted using generalized additive mixed models, with shaded bands indicating 95% confidence intervals. Across the cortex, subcortex and cerebellum, sex-specific trajectories largely recapitulate the dominant age-related patterns observed in the sex-combined analyses, indicating that structure-specific maturation of within-network connectivity is broadly preserved across sexes. Female-male differences are generally modest and appear mainly as differences in trajectory magnitude or late-age divergence in selected systems, rather than qualitatively distinct developmental profiles. These results support the robustness of the main developmental trajectories to sex stratification while indicating limited, network-specific sex modulation across early life.

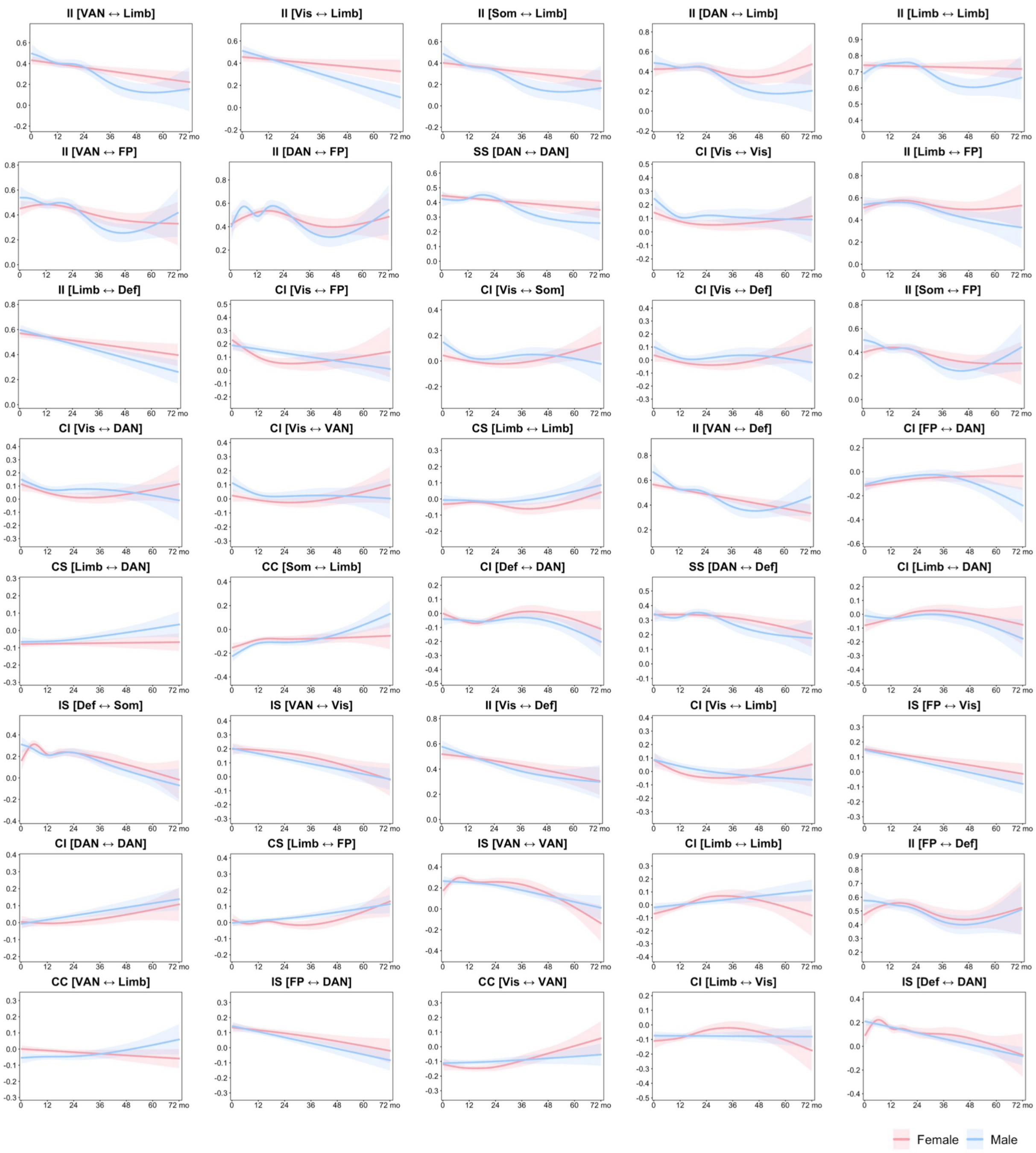

**Supplementary Figure 4**. Developmental trajectories of connections with the largest sex differences. Sex-specific developmental trajectories of functional connectivity are shown for the 40 connections with the largest female-male differences across early life. Connections were ranked by the mean absolute difference between fitted female and male trajectories across the full age range. Trajectories were fitted using generalized additive mixed models, with shaded bands indicating 95% confidence intervals. The displayed connections include both within-structure and cross-structure classes: CC, cortex-cortex; II, cerebellum-cerebellum; SS, subcortex-subcortex; CI, cortex-cerebellum; CS, cortex-subcortex; and IS, cerebellum-subcortex. Network abbreviations: VIS, visual; SMN, somatomotor; DAN, dorsal attention; VAN, ventral attention; LIM, limbic; FPN, frontoparietal; DMN, default mode. Among the displayed connections, sex-related differences were expressed mainly as differences in trajectory magnitude, curvature or late-age divergence, rather than broad reorganization of the developmental connectome.

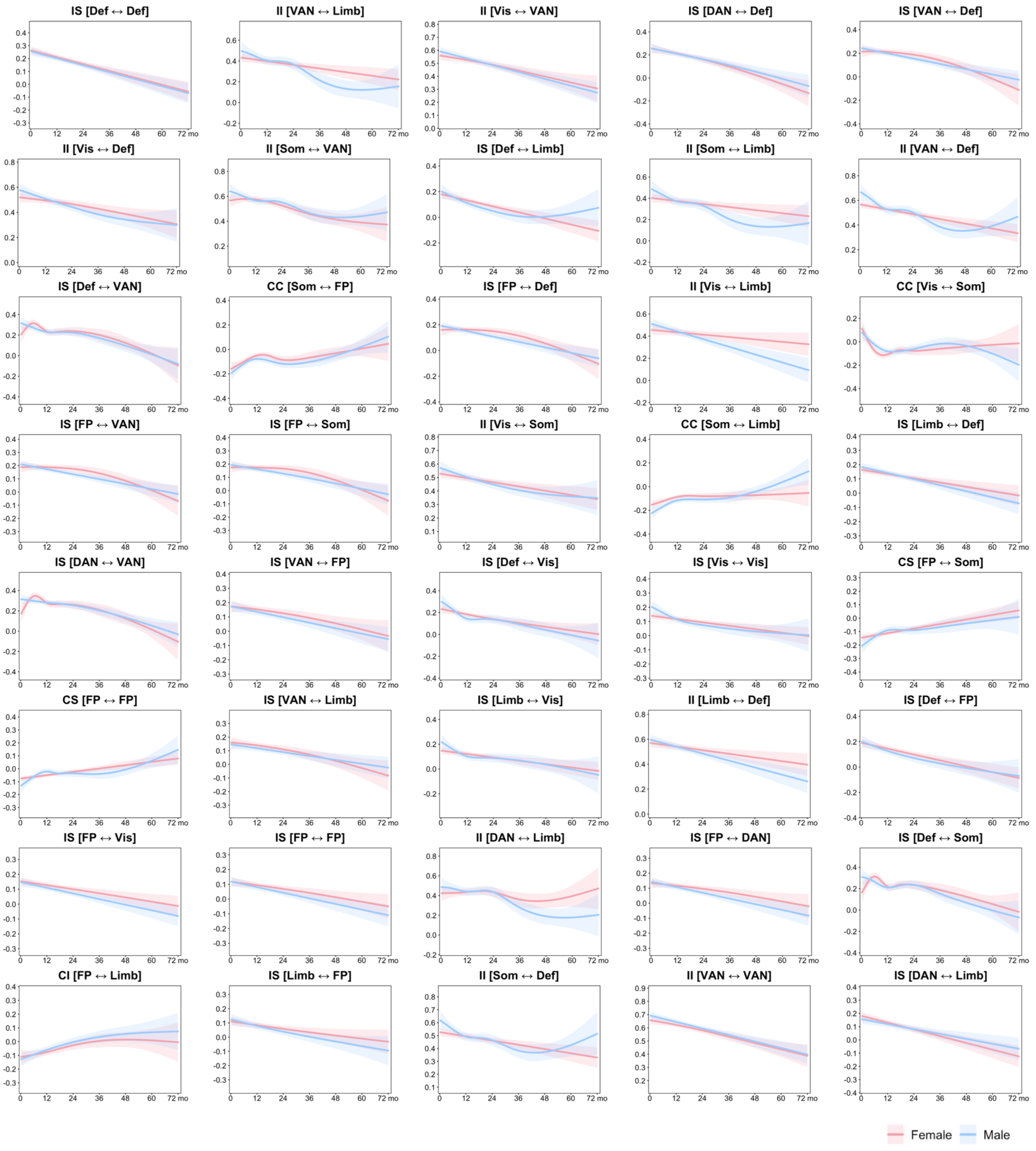

**Supplementary Figure 5**. Developmental trajectories of connections showing the largest age-related differences. Developmental trajectories of functional connectivity are shown for the 40 connections exhibiting the largest age-related differences across early life. Connections were ranked by the absolute difference in fitted connectivity between 0 and 60 months, chosen to avoid sparse late-age boundary estimates, and the top 40 were selected for display. Trajectories were fitted using generalized additive mixed models, with shaded bands indicating 95% confidence intervals. The displayed connections include both within-structure and cross-structure classes: CC, cortex–cortex; II, cerebellum–cerebellum; SS, subcortex-subcortex; CI, cortex-cerebellum; CS, cortex-subcortex; and IS, cerebellum-subcortex. Network abbreviations: VIS, visual; SMN, somatomotor; DAN, dorsal attention; VAN, ventral attention; LIM, limbic; FPN, frontoparietal; DMN, default mode. These trajectories highlight the subset of within- and between-structure connections showing the strongest developmental remodeling across early life.

**Supplementary Tables**

Table S1. Data information

| | BCP | MAP | dHCP | Pixar | Calgary |
|---|---|---|---|---|---|
| **Study design** | Longitudinal | Longitudinal | Cross-sectional | Cross-sectional | Longitudinal |
| **Number of subjects** | 296 | 76 | 88 | 47 | 58 |
| **Total scans** | 597 | 316 | 88 | 47 | 122 |
| **Sex (M/F)** | 139/157 | 37/39 | 62/26 | 20/27 | 31/27 |
| **Age range, months** | 0-73 | 0-76 | 39-45 PMA weeks | 42-72 | 31-83 |

**Table S2. Imaging protocols**

| | | | |
|---|---|---|---|
| **BCP** | ***Anatomical*** | TR/TE (ms) | 2400/2.24 |
| | | Resolution ($mm^3$) | $\mathbf{0.8 \times 0.8 \times 0.8}$ |
| | ***rs-fMRI*** | TR/TE (ms) | 800/37 |
| | | Resolution ($mm^3$) | $\mathbf{2 \times 2 \times 2}$ |
| | | Volumes | 420 |
| | | Imaging state | 0-3 yrs: natural sleep; 3-6 yrs: watching movie |
| **MAP** | ***Anatomical*** | TR/TE (ms) | 1820/4.38 |
| | | Resolution ($mm^3$) | $\mathbf{1 \times 1 \times 1}$ |
| | ***rs-fMRI*** | TR/TE (ms) | 2000/32 |
| | | Resolution ($mm^3$) | $\mathbf{4 \times 4 \times 4}$ |
| | | Volumes | 150 |
| | | Imaging state | 0-2 yrs: natural sleep; 2-6 yrs: watching movie |
| **dHCP** | ***Anatomical*** | TR/TE (ms) | 12000/156 |
| | | Resolution ($mm^3$) | $\mathbf{0.8 \times 0.8 \times 0.8}$ |
| | ***rs-fMRI*** | TR/TE (ms) | 392/38 |
| | | Resolution ($mm^3$) | $\mathbf{2.15 \times 2.15 \times 2.15}$ |
| | | Volumes | 2300 |
| | | Imaging state | Natural sleep |
| **Pixar** | ***Anatomical*** | TR/TE (ms) | 2530/1.64 |
| | | Resolution ($mm^3$) | $\mathbf{1 \times 1 \times 1}$ |
| | ***rs-fMRI*** | TR/TE (ms) | 2000/30 |
| | | Resolution ($mm^3$) | $\mathbf{3 \times 3 \times 3.3}$ |
| | | Volumes | 168 |
| | | Imaging state | Watching movie |
| **Calgary** | ***Anatomical*** | TR/TE (ms) | 8.23/3.76 |
| | | Resolution ($mm^3$) | $\mathbf{0.45 \times 0.45 \times 0.9}$ |
| | ***rs-fMRI*** | TR/TE (ms) | 2000/30 |
| | | Resolution ($mm^3$) | $\mathbf{3.59 \times 3.59 \times 3.6}$ |
| | | Volumes | 250 |
| | | Imaging state | Passive viewing movie |
